# Mucins form a nanoscale material barrier against immune cell attack

**DOI:** 10.1101/2022.01.28.478211

**Authors:** Sangwoo Park, Marshall J. Colville, Carolyn R. Shurer, Ling-Ting Huang, Joe Chin-Hun Kuo, Justin H. Paek, Marc C. Goudge, Jin Su, Matthew P. DeLisa, Jan Lammerding, Warren R. Zipfel, Claudia Fischbach, Heidi L. Reesink, Matthew J. Paszek

## Abstract

The cancer cell glycocalyx serves as a major line of defense against immune surveillance. However, how specific physical properties of the glycocalyx contribute to immune evasion and how these properties are regulated are not well understood. Here, we uncover how the surface density, glycosylation, and crosslinking of cancer-associated mucins contribute to the nanoscale material thickness of the glycocalyx, and further analyze the effect of the glycocalyx thickness on resistance to effector cell attack. Natural Killer (NK) cell-mediated cytotoxicity exhibits a near perfect inverse correlation with the glycocalyx thickness of target cells regardless of the specific glycan structures present. NK cells expressing a chimeric antigen receptor (CAR) have an enhanced ability to breach the glycocalyx and kill target cells. Equipping the NK cell surface with a mucin-digesting enzyme also improves killing with a performance enhancement that rivals or exceeds CARs in some cases. Together, our results provide new considerations for improving cancer immunotherapies.

## INTRODUCTION

Tumor cells construct a glycocalyx with physical and biochemical attributes that guard against detection and destruction by surveilling immune cells(Ghasempour and Freeman, 2021). Essentially all types of human cancers exhibit changes in glycan synthesis, resulting in their expression at atypical levels or with altered structural attributes, phenomena that were first observed more than six decades ago(Hakomori and Murakami, 1967; Kim and Varki, 1997; Ladenson et al., 1949). Immunomodulatory glycans can reprogram the activities of important immune cell types, including macrophages, dendritic cells, cytotoxic T-cells, and Natural Killer (NK) cells(Zhou and Cobb, 2021). Larger macromolecules in the glycocalyx also are proposed to sterically shield molecular epitopes, fundamentally changing how immune cells perceive and interact with tumor cells(Ghasempour and Freeman, 2021). While a multifaceted role for the cancer-associated glycocalyx in immune evasion has become clear, a physical understanding of the glycocalyx remains limited, especially regarding cancer immunotherapy. With the rapid advancement of adoptive cell therapies for cancer, a broader understanding of the function of the glycocalyx in immune evasion has become more urgent.

Cancer cells often generate a glycocalyx with high levels of cell surface mucins, which can directly modulate the tumor immune microenvironment (Burchell et al., 2018; Gupta et al., 2020; Mereiter et al., 2019; RodrÍguez et al., 2018; Wall et al., 2020). Mucin-type O-glycans initiate with N-acetylgalactosamine (GalNAc) that is linked to serine or threonine on the mucin polypeptide backbone and elongated into more complex structures through the sequential actions of glycosyltransferases in the Golgi apparatus(Hanisch, 2001; Narimatsu et al., 2019; Nason et al., 2021). Cancer is often associated with dramatically higher expression of cell-surface mucins that bear truncated and more highly sialylated O-glycans(Burchell et al., 2018; Rømer et al., 2021). Through the engagement of Siglecs, the macrophage galactose receptor, and other immune-cell receptors, tumor-associated O-glycans have been implicated in the direct suppression of CD4^+^ T, CD8^+^ T, and NK cell activities(Hudak et al., 2014a; Perdicchio et al., 2016; Wall et al., 2020), as well as the recruitment and education of tumor-associated macrophages(Beatson et al., 2016; Dusosw et al., 2020; Rodriguez et al., 2021). Mucins and cancer-associated O-glycans also serve as scaffolds for multivalent galectins that can act as negative regulators, or checkpoints, of the immune cell functions(Compagno et al., 2020; Liu and Rabinovich, 2005; Perillo et al., 1995). Immune suppression by mucins and mucin-type O-glycans is under active investigation as a mechanism of recalcitrance to cancer immunotherapies.

From a physical perspective, cell-surface mucins are semiflexible biopolymers that form a soft material coating on the tumor-cell surface(Kuo et al., 2018; Paszek et al., 2014). Although recent studies of immune modulation by mucins have focused on the biochemical activity of their O-glycans, it has been suggested that the dense grafting of mucin biopolymers on the tumor cell surface may form a steric barrier against immune-cell recognition and cytolytic activities(Hollingsworth and Swanson, 2004; Madsen et al., 2013; Suzuki et al., 2012). Direct validation of this possibility has remained elusive due to the challenge of disentangling the biochemical and material effects of specific changes in mucin-type glycosylation(Madsen et al., 2013). The dense clustering of O-glycans along the mucin backbone both extends and rigidifies its macromolecular structure(Hanisch, 2001; Kramer et al., 2015; Kuo et al., 2018). Therefore, changes in the number, length, and sialylation (i.e. charge) of O-glycan chains are expected to alter the molecular properties of individual mucins and the ensemble material properties of the mucin-rich glycocalyx(Kuo et al., 2018). Given the close, intimate contact that is required for target engagement by effector immune cells, the material thickness of the glycocalyx may be a major determinant of immune evasion. However, the investigation of how the physical dimensions of the glycocalyx change in response to altered states of glycosylation has been challenging due the nanometer-scale precision of measurement that is required to detect the structural changes(Kuo and Paszek, 2021).

Clinical interest in Natural killer (NK) cells for adoptive cell therapy has risen swiftly due to their ability to kill cells bearing markers associated with oncogenic transformation(Shimasaki et al., 2020). NK cells have been equipped with chimeric antigen receptors (CAR-NK) to direct attacks against specific tumor antigens, with some CAR-NK therapeutics now in human clinical trials(Liu et al., 2020; Shimasaki et al., 2020; Tang et al., 2018; Xiao et al., 2019). Compared to CAR-T cells, CAR-NK cells have some significant advantages that may include better safety and more potential for “off-the-shelf” manufacturing of allogeneic therapies. As is the case with other immune therapies, tumors can develop multiple mechanisms to resist attack by NK and CAR-NK cells. Notably, an inverse relationship between NK cell killing and the expression of cell-surface mucins on target cells has been reported(Madsen et al., 2013). Truncation of mucin-type O-glycans increases NK cell-mediated antibody-dependent cellular-cytotoxicity, whereas elongation of the O-glycans has been reported to protect tumor cells(Madsen et al., 2013; Okamoto et al., 2013; Suzuki et al., 2012; Tsuboi et al., 2011). How such changes alter the material properties of the glycocalyx is unknown, and it remains unclear if the effects on NK-cell resistance are caused by specific receptor interactions, changes in the structural attributes of the mucins, or some other consequence of altered glycosylation(Madsen et al., 2013). There remains a need for new tools that can probe and measure the material properties of the glycocalyx to understand how these properties may contribute to immune evasion.

Recently, several imaging strategies have been developed to image the nanoscale dimensions and structural organization of the glycocalyx(Kuo and Paszek, 2021; Möckl et al., 2019; Paszek et al., 2012; Son et al., 2020). One of these technologies, called Scanning Angle Interference Microscopy (SAIM), is a localization microscopy technique based on the principles of Fluorescent Interference Contrast Microscopy (FLIC) whereby standing waves of excitation light are used to axially localize fluorescently labelled structures of interest with nanoscale precision(Ajo-Franklin et al., 2005; Carbone et al., 2016; Colville et al., 2019; Lambacher and Fromherz, 1996; Paszek et al., 2014). To improve the precision of SAIM for glycocalyx materials research, we recently reported a new implementation that uses a pair of high-speed, galvanometer-controlled mirrors to generate a revolving circle, or “ring”, of excitation light at defined sample incidence angles(Colville et al., 2019). The approach, which we refer to here as Ring-SAIM, is intended to improve sample illumination and, thus, the precision of measurement compared to standard SAIM implementations (Colville et al., 2019).

Taking advantage of Ring-SAIM, we demonstrate that the material thickness of the glycocalyx is an important parameter that can explain cellular resistance to NK and CAR-NK attack. By combining approaches for nanoscale optical imaging with genetic approaches to edit the molecular composition of the glycocalyx, we show how specific molecular features of mucins contribute to the material thickness of the glycocalyx. Our results uncover new strategies, including the surface display of glycocalyx editing enzymes, to penetrate the glycocalyx barrier for improved cytotoxicity by immune cells.

## RESULTS

### Physical properties of the mucin polymer brush

Cell-surface mucins are anchored to the plasma membrane on one end by their transmembrane domain, forming a brush-like structure(Button et al., 2012; Shurer et al., 2019; Song et al., 2020). Polymer chains are predicted to extend out at higher surface densities due to steric repulsions in a more crowded structure(Milner, 1991). In tumors and cell lines that overexpress cancer-associated mucins, high heterogeneity in the cell surface levels is typical(O’Connor et al., 2005; Walsh et al., 1999). To test how the expression level of mucin affects the material thickness of the glycocalyx, we developed a cellular model with highly titratable cell-surface levels of the cancer-associated mucin, Muc1. We constructed an expression cassette for Muc1-GFP with 42 TRs under the control of a tetracycline-inducible promoter (Figure 1A). We initially generated a polyclonal cell population through stable integration of the expression cassette in epithelial cells. In the polyclonal population, our ability to titrate mucin expression in a dose-dependent manner with doxycycline was limited by the large cell-to-cell variability in expression levels and the variability of the doxycycline response curves. To overcome this challenge, we clonally expanded the cell pool and identified multiple clones that exhibited ideal dose responses to doxycycline induction. One of these clonal lines, referred to here as 1E7, was selected as our model for subsequent experiments (Figure 1B-1D).

**Figure 1.**
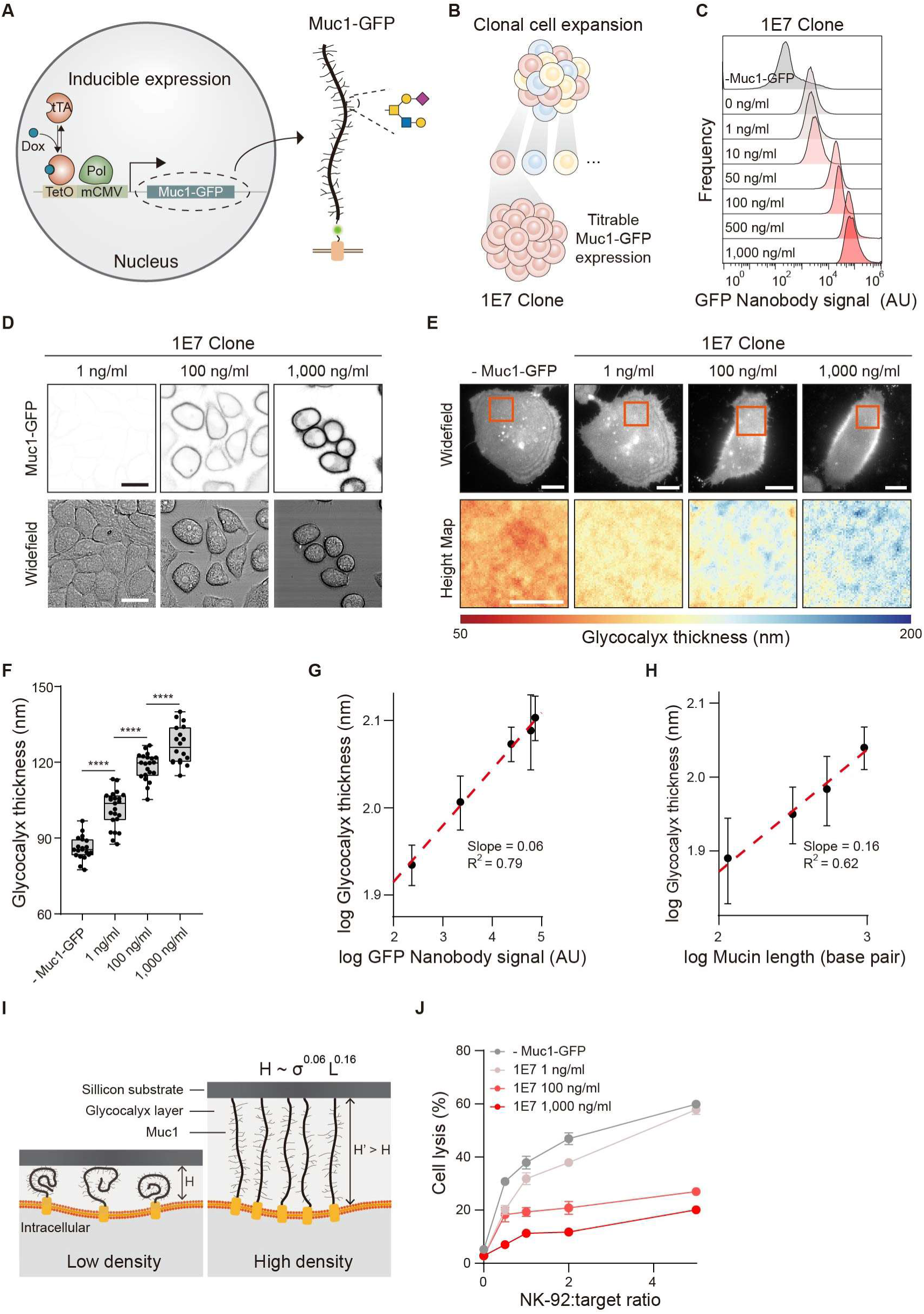
Physical behavior of the mucin material barrier. (A) Doxycycline inducible system for controlled expression of Muc1-GFP with 42 tandem repeats. (B) Cartoon showing clonal expansion of epithelial cells expressing Muc1-GFP to isolate a stable clone, 1E7, with titratable mucin expression levels. (C) Flow cytometry analysis of the 1E7 clone showing titratable Muc1-GFP levels across the indicated concentrations of doxycycline inducer. (D) Fluorescence and widefield images of 1E7 cells induced at the indicated concentration of doxycycline (Scale bar, 10 µm). (E) Representative widefield fluorescence images and Ring-SAIM reconstructions of glycocalyx thickness in 1E7 cells at the indicated doxycycline induction levels and control epithelial cells that do not express Muc1-GFP (-Muc1-GFP). (F) Quantification of mean glycocalyx thickness in control cells (-Muc1-GFP) and 1E7 cells at the indicated induction levels. Results are the mean ± s.d. of at least 18 cells. Statistical significance is given by **** P ≤ 0.0001. (G) Log of the mean glycocalyx thickness in 1E7 cells versus the log of the mean GFP-nanobody signal, a proportionate measure of Muc1 surface density; best-fit line to a power-law model has slope = 0.06 and *R*^2^ = 0.79. (H) Log of the mean glycocalyx thickness versus log of the mucin biopolymer length; best-fit line to a power-law model with slope = 0.16 and *R*^2^ = 0.62. (I) Schematic summarizing the scaling behavior of the glycocalyx layer on mucin density, σ, and chain size, L. (J) NK-92 cell-mediated lysis of control cells (-Muc1-GFP) and 1E7 target cells at the indicated doxycycline induction levels; results are shown for varying NK-92 to target cell ratios. Results are a representative dataset from 3 independent replicates. (Scale bar, 10 µm).

Noting that the height and compressibility of a polymer brush are known to scale with the polymer grafting density and polymer chain length(Gandhi et al., 2019; Gennes, 1987), we sought to develop quantitative relationships that described the thickening of the glycocalyx by increased expression of cell-surface mucins. We first validated that our Ring-SAIM setup could detect nanometer-scale changes in the glycocalyx thickness of live cells using Muc1 constructs of varying size (Figure S1). After confirming the ability of Ring-SAIM to detect changes in glycocalyx thickness of approximately 10-nm (Figure S1H), we measured the thickness of the glycocalyx in 1E7 cells at varying doxycycline induction levels. We observed a progressive thickening of the glycocalyx with higher induction levels of Muc1 (Figure 1E). The glycocalyx thickness ranged from 86.0 ± 4.58 nm for cells without Muc1-GFP expression to 126.8 ± 7.45 nm for cells at saturating levels of Muc1-GFP induction (Figure 1F). Our data revealed a clear power-law dependence of glycocalyx thickness on the Muc1-GFP surface density with a scaling exponent of 0.06 (Figure 1G). To construct this relationship, we used the GFP nanobody signal as a proportionate quantity to the Muc1-GFP surface level. The glycocalyx thickness also had a clear power-law dependence on the length of the mucin with a scaling exponent of approximately 0.16 (Figure 1H). Thus, the glycocalyx thickness weakly scaled with mucin surface density and more strongly scaled with mucin backbone length (Figure 1I). These relationships provide an important framework for understanding how the heterogeneity in mucin expression level and variation in the number of TRs could influence the thickness and barrier properties of the glycocalyx (Figure 1I).

### The mucin brush resists NK-cell attack

The SAIM measurements in the 1E7 clone indicated how mucin chain length and density would resist close cellular contact with a surface (Figure 1G and 1H). We tested whether a mucin brush could similarly resist effector cell engagement with a target cell, a process that requires close, receptor-mediated interactions between cells. We considered immune cell attack by NK-92, a well-characterized NK cell line that has demonstrated promising anticancer activities in human clinical trials(Arai et al., 2008; Williams et al., 2017). NK-92 cytotoxicity against target cells was reduced by Muc1-GFP expression in a dose-dependent manner (Figure 1J). NK-92 cells do not express appreciable levels of inhibitory Siglec-7 and Siglec-9, the two primary receptors which have been implicated in NK suppression by Muc1 O-glycans(Hudak et al., 2014b; Rosenstock et al., 2017; Wall et al., 2020), raising the possibility that the protection by mucins was mediated by a physical mechanism. To ensure that the results were not a clonal artifact limited to the 1E7 cell line, we repeated the NK-92 cytotoxicity assay against two additional Muc1-GFP expressing clones, as well as the parental polyclonal cell line from which the 1E7 clone was derived. We observed a similar trend of enhanced resistance to NK-cell attack with increasing Muc1-GFP expression in the parental line and all clones tested (Figure S2).

### Regulation of glycocalyx material thickness by mucin-type O-glycosylation and multivalent galectins

We sought to develop an understanding of how specific molecular attributes of mucins might contribute to the ensemble materials properties of the glycocalyx and, thus, protection against immune cell attack. For bottlebrush copolymers with densely grafted side chains, such as mucins, the size and charge of the side chains can strongly influence the polymer rigidity and persistence length(Paturej et al., 2016; Sarapas et al., 2020). Thus, we investigated how O-glycan truncation, elongation, and sialylation each contribute to the material thickness of the glycocalyx in 1E7 cells.

O-glycans were truncated in 1E7 cells through CRISPR/Cas9 mediated knockout (KO) of the *C1GALT1* gene, which encodes the Core 1 synthase that governs the extension of α-GalNAc (Figure 2A). Sialylation was ablated by CRISPR/Cas9 KO of the *GNE* gene, which encodes the glucosamine (UDP-N-acetyl)-2-epimerase/N-acetylmannosamine kinase, an essential enzyme in CMP-sialic acid biosynthesis. The 1E7 *C1GALT1* and *GNE* KO cells were each clonally expanded to isolate homozygous KO lines (Figure S3A and S3B). Glucosaminyl (N-Acetyl) Transferase 1 (GCNT1) was overexpressed to increase O-glycan extension through Core-II elongation (Figure 2A and S3C). Analysis by flow cytometry showed similar cell-surface levels of Muc1-GFP across the glycoengineered cell line panel, indicating that manipulation of glycosylation did not alter Muc1-GFP surface expression (Figure 2B). Disruption of Core-I O-glycan extension or sialylation in the KO lines was confirmed by flow cytometry using appropriate lectin probes (Figure 2B). Since there are no direct immunochemical or lectin probes for Core-II O-glycans, we instead confirmed increased elongation of O-glycans in GCNT1 overexpressing cells by SDS-PAGE (Figure 2C and 2D). The apparent molecular weight of Muc1-GFP observed by SDS-PAGE was substantially lower in *C1GALT1* KO cells and appreciably higher in GCNT1 overexpressing cells compared to Muc1-GFP in wild-type cells, consistent with the presence of shorter and longer glycan side chains, respectively (Figure 2C and 2D).

**Figure 2.**
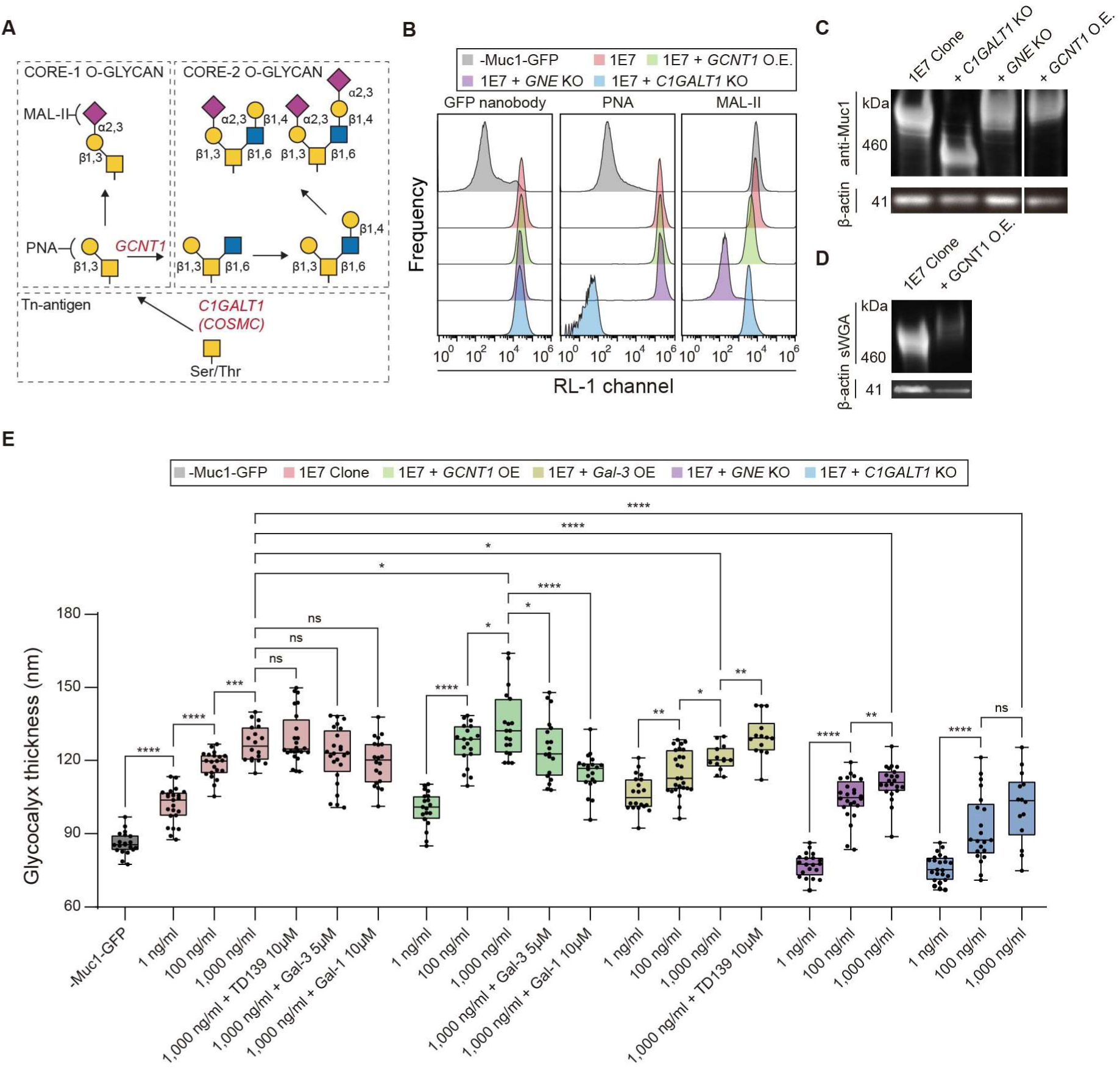
Molecular determinants of the mucin material barrier. (A) Biosynthetic pathway and lectin probes for mucin-type core-I and core-II O-glycans. (B) Flow cytometry analysis of 1E7 cells and their glycoengineered progeny with knockout (KO) of *C1GALT1* to truncate O-glycans, KO of *GNE* to ablate glycan sialylation, and overexpression (OE) of *GCNT1* to enhance O-glycan extension; results for epithelial cells that do not express Muc1-GFP (-Muc1-GFP) are included as a control. (C) Western blot showing the relative size of Muc1 polymer in the 1E7 clone and glycoengineered progeny. (D) sWGA lectin blot after extended SDS-PAGE run time showing increased molecular weight of Muc1-GFP in GCNT1-overexpressing 1E7 cells compared to wild-type 1E7 cells. (E) Quantification of glycocalyx thickness in 1E7 cells, their glycoengineered progeny, and galectin-3 overexpressing cells (+Gal3 OE) at the indicated doxycycline induction levels; as indicated, some treatments include 10 µM recombinant galectin-1, 5 µM recombinant galectin-3, or 10 µM of the galectin inhibitor TD139. Results are the mean ± s.d. of at least 13 cells. Statistical significance is given by ns - not significant, * P ≤ 0.05, ** P ≤ 0.01, *** P ≤ 0.001, **** P ≤ 0.0001. Results are a representative dataset from 3 independent replicates.

Utilizing Ring-SAIM, we measured the material thickness of the glycocalyx across the glycoengineered cell panel. At each Muc1-GFP induction level, the glycocalyx was significantly thinner in *C1GALT1* and GNE KO cells compared to wild-type 1E7 cells (Figure 2E). At 1,000 ng/mL doxycycline induction, KO of *C1GALT1* had the largest effect on the glycocalyx, reducing its material thickness by 25.1 nm compared to wild-type 1E7 cells. KO of GNE reduced the average glycocalyx thickness by 16 nm, whereas GCNT1 overexpression increased the average glycocalyx thickness by 7.6 nm at the same induction level (Figure 2E). Importantly, even with the truncation of O-glycans or ablation of sialylation, the thickness of the glycocalyx increased with increasing surface density, as expected for a polymer brush (Figure 2E). Thus, even minimally glycosylated mucin polymers could resist close cell-to-surface contact when expressed at sufficiently high levels.

Multivalent galectins have been proposed to crosslink and increase the barrier properties of cell-surface mucins(Argüeso et al., 2009). Consistent with prior observations(Earl et al., 2010; Yu et al., 2007; Zhao et al., 2009), we confirmed specific interactions of galectin-1 and -3 with Muc1-GFP on 1E7 cells (Figure S4). Recombinant galectin-1 showed a preference for Core-II O-glycans, whereas recombinant galectin-3 bound specifically to Core-I O-glycan structures, as well as N-glycans (Figure S4). Treatment of wild-type 1E7 cells with recombinant galectin-3, recombinant galectin-1, or 10 µm of the small-molecule galectin inhibitor TD139 (K_d_ = 0.036 µM for galectin-3; K_d_ = 2.2 µM for galectin-1) did not significantly change the material thickness of the glycocalyx at 1,000 ng/mL doxycycline induction (Figure 2E). Likewise, the effect of galectin-3 overexpression on the glycocalyx thickness in 1E7 cells was either small or not statistically significant depending on the Muc1-GFP expression level (Figure 2E). In 1E7 cells overexpressing GCNT1, treatment with recombinant galectin-3 slightly decreased the glycocalyx thickness by 9.5 nm compared to untreated cells (p = 0.0308; Figure 2E). Galectin-1 treatment had a more pronounced effect, reducing the glycocalyx thickness by 19.4 nm compared to untreated GCNT1 overexpressing cells (p = 0.000003; Figure 2E).

These observations were consistent with galectins restricting the free chain extension of mucins, presumably through either intermolecular or intramolecular crosslinking. Fluorescence recovery after photobleaching (FRAP) showed that a large percentage of mucins were mobile in the cell-surface brush, indicating that intermolecular crosslinking by galectin was modest (Figure S5). Overall, the relatively small effects of the galectin manipulations compared to the larger effects observed with varying mucin surface density, length, and glycosylation pointed to chain entropy, and not crosslinking, as the major determinants of the glycocalyx material thickness at the substrate interface.

### The material thickness of the glycocalyx accounts for the resistance to NK cell attack

Our Ring-SAIM measurements and biochemical analyses provided a roadmap for how the material and biochemical state of the glycocalyx could be manipulated by tuning mucin density and glycosylation. Armed with this information, we next tested how specific glycocalyx states would protect against immune cell attack. 1E7 cells at varying Muc1-GFP induction levels and their glycoengineered progenies were co-incubated with NK-92 to assess cytotoxicity. Across the glycoengineered panel, NK-92 mediated cytotoxicity against the mucin-expressing target cells was progressively reduced at higher doxycycline induction levels (Figure 3A). Mucins with truncated and less sialylated O-glycans provided less protection against NK-92 attack (Figure 3A). Thus, resistance to NK cell attack appeared to correlate with conditions that favored extension of a thicker glycocalyx.

**Figure 3.**
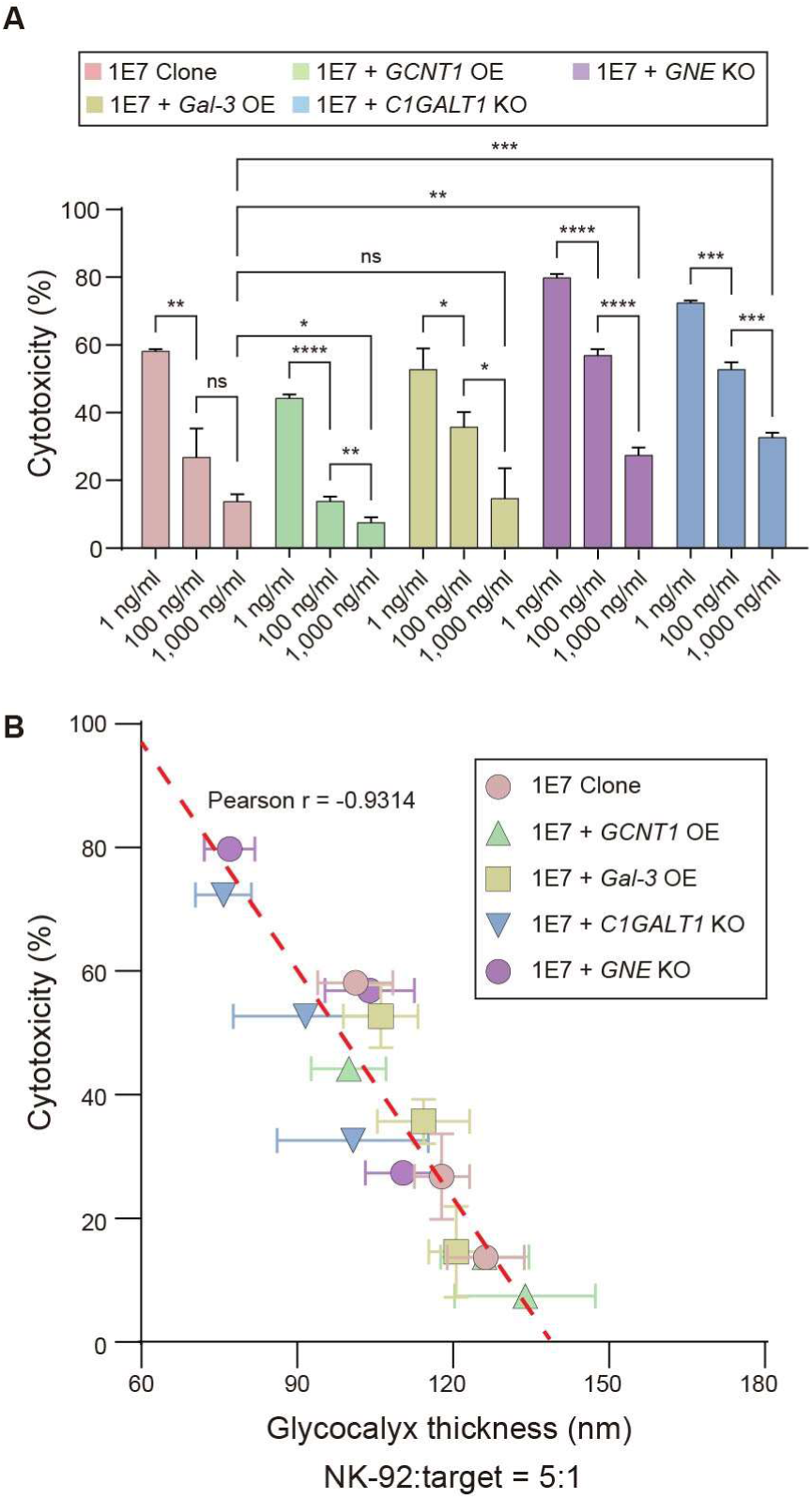
The mucin material barrier guards against NK cell-mediated cytotoxicity. (A) NK-92 cell-mediated cytotoxicity of 1E7 cells, their glycoengineered progeny (*C1GALT1* KO, *GNE* KO, and GCNT1 OE), and 1E7 cells overexpressing galectin-3 (Gal-3 OE) at the indicated doxycycline induction level; NK cell to target cell ratio is 5:1. (B) NK-92 cell-mediated cytotoxicity is inversely proportional to the measured glycocalyx thickness with Pearson correlation coefficient, r = -0.94; dashed line indicates a linear fit to the data. Data presented as the mean and ± s.d. of three independent experiments. Statistical significance is given by ns - not significant, * P ≤ 0.05, ** P ≤ 0.01, *** P ≤ 0.001, **** P ≤ 0.0001. Results are a representative dataset from 3 independent replicates.

To assess the relationship between resistance to immune cell attack and glycocalyx material thickness, the NK-mediated cytotoxicity reported in Figure 3A was plotted against the measured glycocalyx thickness reported in Figure 2E for each of the glycoengineered cell type and Muc1-GFP induction level. We observed a strong and nearly perfect inverse correlation between cytotoxicity and glycocalyx material thickness across the combined dataset representing different mucin expression levels and glycosylation patterns (Pearson coefficient = -0.94; Figure 3B). The cytotoxicity data were fit to a linear model dependent on a single variable: the glycocalyx thickness (Figure 3B). Remarkably, we found that the glycocalyx thickness explained 86% of the variation in NK mediated cytotoxicity against the full glycoengineered cell panel, which presents glycocalyces with widely varying glycosylation patterns and biochemical attributes. These results strongly implicate the glycocalyx material structure as a major determinant of cellular resistance to immune cell attack.

### Tethering mucinase to NK cells can dramatically increase their cytotoxicity against mucin-expressing target cells

Enterohemorrhagic *Escherichia coli* (EHEC) utilize the mucin-specific metalloprotease, StcE, to breach the gut mucosal barrier and adhere to intestinal cells(Yu et al., 2012). We reasoned that the StcE mucinase might similarly be equipped on effector immune cells to breach the mucin barrier on target cells. Indeed, the StcE mucinase was highly effective at removing the cell-surface mucin brush from 1E7 cells (Figure S6A and S6B), rendering the cells more susceptible to NK-92 mediated attack (Figure S6C). Motivated by this finding, we investigated how the StcE mucinase could be tethered to the NK-92 cell surface. Noting that the StcE mucinase possesses a C-terminal lectin domain, referred to as X409, in addition to its catalytic domain(Nason et al., 2021), we tested whether the StcE mucinase would specifically bind to the NK-92 cell surface and remain bound without an additional molecular tether. The recombinant StcE bound with high affinity to the NK-92 cell surface and was retained on the NK-92 surface for prolonged periods even after stringent washing with buffer (Figure 4A and 4B). StcE-ΔX409 with a deleted C-terminal lectin domain was not retained on the NK-92 surface, indicating that the X409 domain is the primary binding module to the NK-92 surface (Figure 4B).

**Figure 4.**
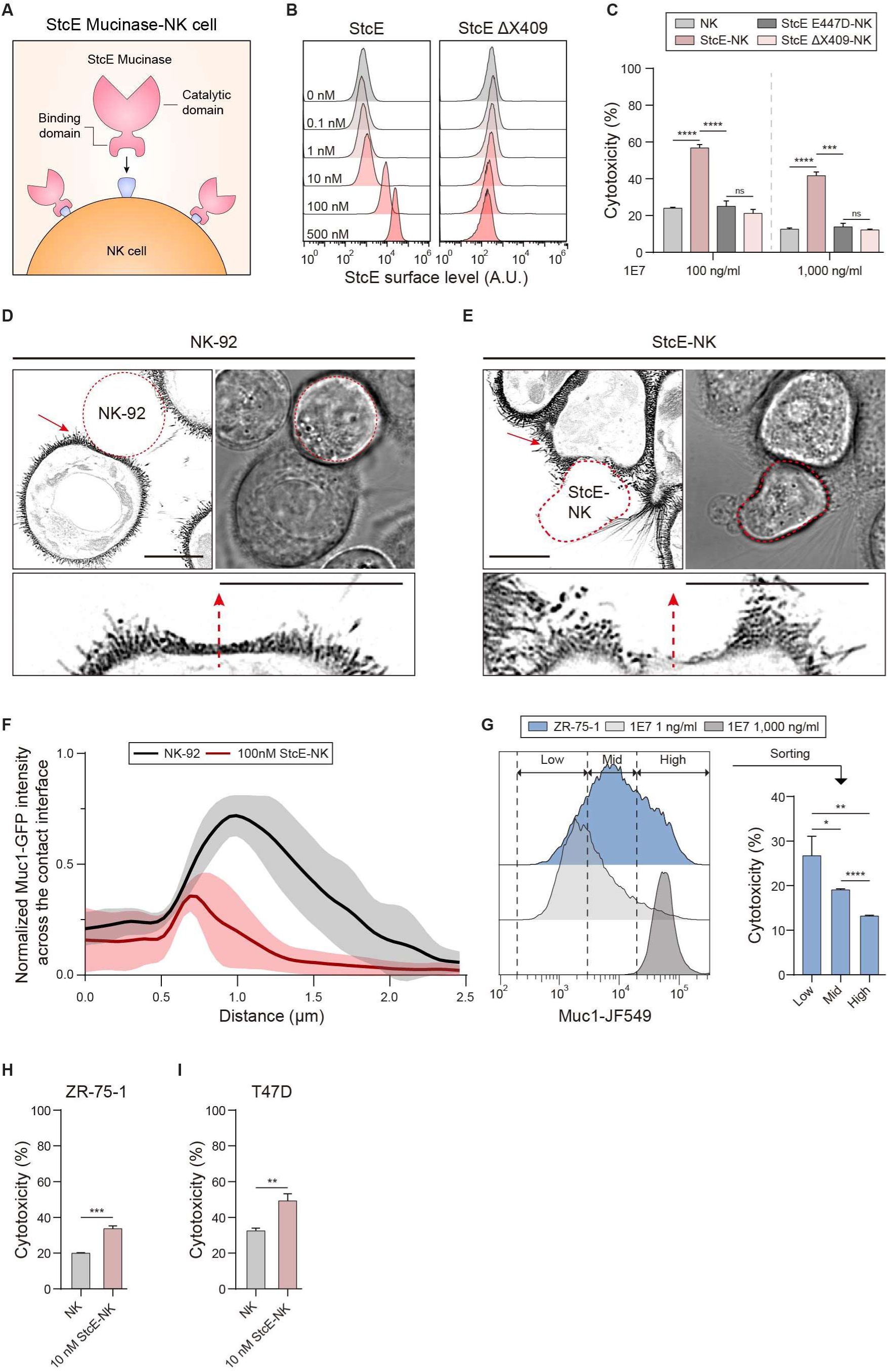
Tethering StcE mucinase to the NK cell surface enhances their cytolytic efficiency. (A) Cartoon showing preparation of StcE-NK through tethering of StcE mucinase to the NK-92 cell surface by its X409 lectin domain. (B) Flow cytometry analysis of StcE surface levels on NK cells incubated with wild-type StcE and StcE with deletion of the X409 lectin domain (StcE ΔX409) at the indicated concentration; StcE surface level probed with anti His-tag antibody. (C) Cytotoxicity of NK-92 cells, StcE-NK cells, StcE E447D-NK cells, and StcE ΔX409-NK cells against 1E7 cells at the indicated doxycycline induction level of Muc1-GFP expression; StcE-NK cells, StcE E447D-NK cells, and StcE ΔX409-NK were prepared through incubation with 100 nM StcE, catalytically-dead StcE E447D, or StcE ΔX409; NK cell to target cell ratio is 10:1. (D, E) Representative Muc1-GFP and brightfield images of 1E7 cells interacting with NK-92 (D) and StcE-NK (E) cells; lower images show zoomed-in regions of the NK-1E7 interface (Scale bar, 10 µm). (F) Plot of the mean Muc1-GFP intensity (dark lines) and standard deviations (lighter bounding areas) along line profiles across the contact interface between 1E7 cell and NK cells (n=6) or StcE-NK cells (n=10); the fluorescence intensity is normalized to the mean fluorescence intensity outside of the contact interface; dotted arrows in D and E show representative profile lines. (G) Cytotoxicity of NK cells against ZR-75-1 breast cancer cells sorted into three sub-populations with low, medium, and high levels of endogenous Muc1expression; NK cell to target cell ratio is 5:1. (H, I) Cytotoxicity of NK cells and 10nM StcE-NK cells against unsorted ZR-75-1 (H) and T47D (I) breast cancer cells that endogenously express Muc1; NK cell to target cell ratio is 5:1. Statistical significance is given by * P ≤ 0.05, ** P ≤ 0.01, *** P ≤ 0.001, **** P ≤ 0.0001.

The StcE-equipped NK-92 (StcE-NK) cells were considerably more effective at killing mucin-expressing target cells than the unmodified NK-92 cells (Figure 4C). Control experiments with the catalytically dead StcE-E447D mucinase confirmed that the enzymatic activity of StcE was required for the enhancement of cytotoxicity (Figure 4C). Additional controls indicated that the surface tethering of StcE to the NK surface was required for the enhanced cytotoxicity against mucin-expressing targets. Indeed, the cytolytic activity of NK cells was not significantly enhanced by treatment with StcE-ΔX409, which we confirmed could actively digest Muc1 but did not remain tethered to the NK surface (Figure 4C and S6A). The NK-92 cells possessed a small number of mucin-domain containing proteins that were trimmed similarly by wild-type StcE and StcE-ΔX409 (Figure S6D). The inability of StcE-ΔX409 treatment to enhance the cytolytic activity of NK cells indicated that the StcE mucinase treatment did not directly enhance the latent activity of NK cells through the trimming of mucin domain proteins from the NK cells surface (Figure 6C).

We observed a localized disruption of the mucin barrier at the interface between StcE-NK cells and target cells. Visible deflection of microvilli on 1E7 cells in the contact zone with both NK-92 and StcE-NK indicated the formation of viable mechanical connections (Figure 4D and 4E). However, a high density of Muc1-GFP and microvilli persisted in the contact zone with unmodified NK-92 cells (Figure 4D). Significantly lower Muc1-GFP levels and fewer microvilli were observed in the contact zone with StcE-NK cells (Figure 4E and 4F). These results were consistent with the surface-tethered StcE locally cleaving mucin or otherwise aiding in the segregation of cell-surface mucin from the contact interface, as well as disrupting microvilli (Figure 4F).

We next tested the ability of StcE-NK cells to attack cancer cell lines that endogenously express cell-surface mucins. We benchmarked the Muc1 surface levels of a luminal A breast cancer cell line, ZR-75-1, against the 1E7 cell line. Non-permeabilized cells were probed with conventional Muc1 monoclonal antibodies and analyzed by flow cytometry. We constructed distributions for Muc1 surface levels in 1E7 cells at minimal and saturating Muc1-GFP induction (1 ng/mL and 1,000 ng/mL doxycycline) to define standardized gates for low, moderate, and high Muc1 expression that could be applied to categorize other cell lines. The Muc1 surface levels were highly heterogeneous in the ZR-75-1 cell line and fully overlapped with the 1E7 low, medium, and high expression benchmarks (Figure 4G). Notably, these results confirmed that the Muc1-GFP levels investigated in the model 1E7 cell system were within the bounds of naturally varying Muc1 levels in tumor cells.

The resistance of NK-92 mediated attack increased progressively with higher cell-surface Muc1 levels on the target ZR-75-1 cells (Figure 4G). StcE-NK cells attacked the ZR-75-1 cells more aggressively compared to unmodified NK-92 cells (Figure 4H). Similar results were observed in T47D cells, another luminal A breast cancer line that expresses moderate to high levels of endogenous Muc1 (Figure 4I). Together, our results suggested that StcE display could improve the efficacy of killer cell attack against targets that are protected by a mucin barrier.

### CAR-NK and StcE-NK can overcome the mucin barrier

Chimeric antigen receptors (CAR) engagement in CAR-NK cells can override inhibitory signals deployed by tumor cells and directly trigger the effector cells’ intrinsic cytolytic effector functions, as well as the release of pro-inflammatory cytokines(Zhang et al., 2017). We next tested whether a mucin barrier would also protect against CAR-NK attack. We engineered the NK-92 cell line to express a humanized anti-HER2 CAR with CD28 and CD3ζ signaling domains (Figure 5A). This CAR was previously validated in NK-92 cells, which are currently under investigation in human clinical trials for glioblastoma(Schönfeld et al., 2015; Zhang et al., 2014).

**Figure 5.**
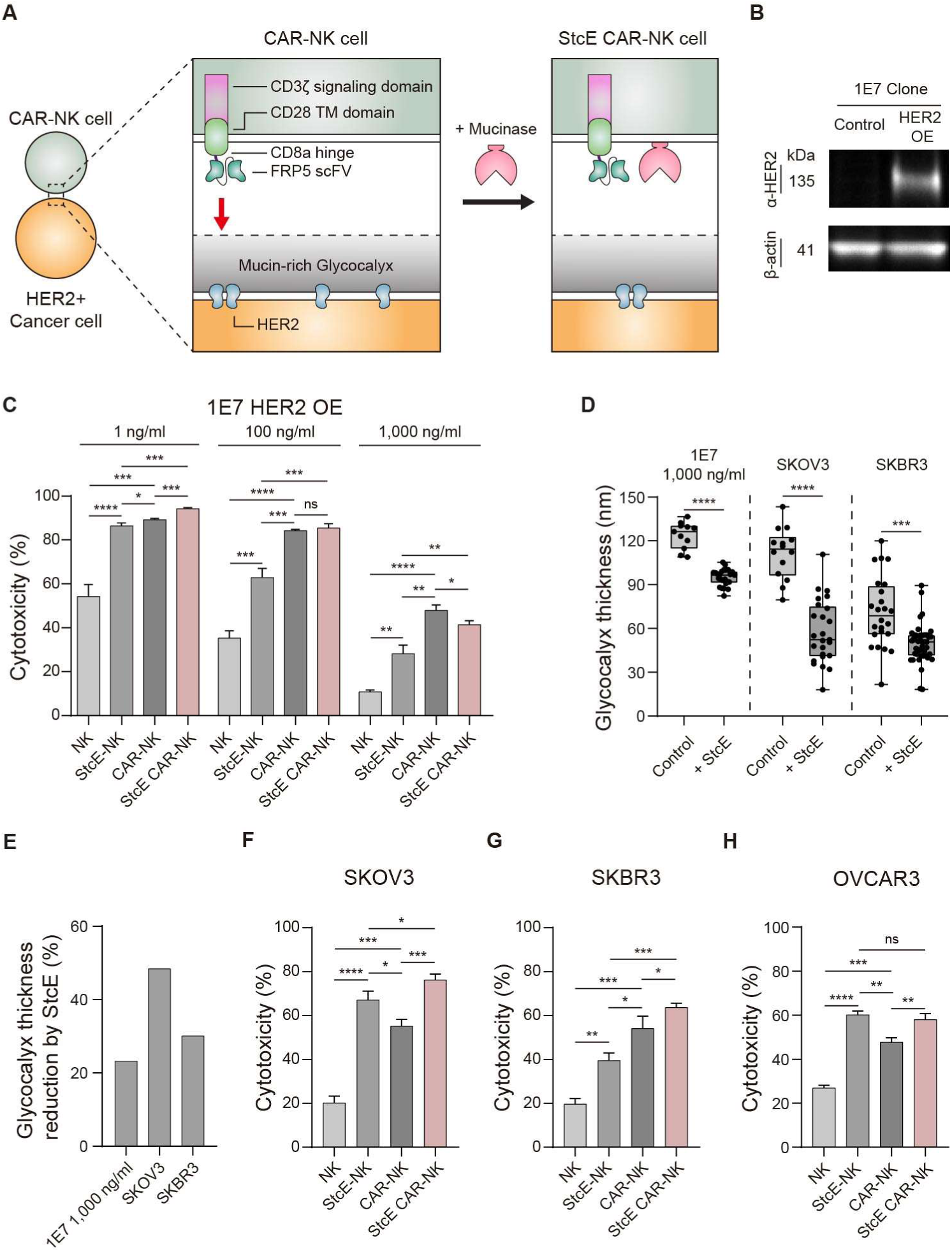
The glycocalyx is a barrier against engineered immune cell therapies. (A) Cartoon showing construction of NK-92 cells equipped with HER2 chimeric antigen receptor (CAR-NK) and CAR-NK cells additionally equipped with StcE mucinase (StcE CAR-NK). (B) Western blot analysis of wild-type (control) and HER2 overexpressing 1E7 cells. (C) Cytotoxicity of NK, StcE-NK, CAR-NK, and StcE CAR-NK against HER2-overexpressed 1E7 clone at the indicated concentration of doxycycline; NK cell to target cell ratio is 5:1. (D) Quantification of mean glycocalyx thickness in the indicated cells before and after treatment of StcE; 1E7 cells induced 1,000 ng/ml doxycycline. (E) Percent reduction of glycocalyx thickness by StcE treatment in the indicated cells from (D). (F, G, H) Cytotoxicity of NK, StcE-NK, CAR-NK, and StcE CAR-NK against SKOV3 (F), SKBR3 (G) and OVCAR3 (H). NK cell to target cell ratio is 10:1. Statistical significance is given by * P ≤0.05, ** P ≤ 0.01, *** P ≤ 0.001, **** P ≤ 0.0001.

To test the ability of the glycocalyx to physically resist CAR-NK cell attack, HER2 was overexpressed in the 1E7 cell line (Figure 5B). CAR-NK cells were more effective at killing HER2 1E7 cells than control NK-92 cells, although the mucin barrier still provided resistance against CAR-NK attack (Figure 5C). Cytotoxicity mediated by CAR-NK cells against HER2 1E7 cells dropped from 89.40% at 1 ng/mL doxycycline induction of Muc1-GFP to 47.92% at 1,000 ng/mL induction (Figure 5C). CAR-NK cells killed the HER2 1E7 targets with moderately higher efficiency compared to StcE-NK cells (Figure 5C). StcE CAR-NK cells, which were equipped with mucinase and the engineered immune receptor, killed the 1E7 targets with similar efficiency as CAR-NK cells (Figure 5A and 5C).

We evaluated the killing efficiency of StcE-NK, CAR-NK, and StcE CAR-NK cells against cancer cells lines that endogenously express HER2 and cell-surface mucin. We first tested the ovarian cancer cell line, SKOV3, which expresses Muc1 and another large mucin called Podocalyxin(Cipollone et al., 2012; Wang et al., 2011). SKOV3 cells had a thick glycocalyx of approximately 120 nm that was reduced to approximately 60 nm following mucinase treatment, confirming that cell-surface mucins were predominant structural elements of the glycocalyx of these cells (Figure 5D). StcE-NK cells had more than three-fold higher cytotoxicity against SKOV3 targets than unmodified NK-92 cells, killing the targets with an even greater efficiency than CAR-NK cells (Figure 5F). We also tested SKBR3 breast cancer cells, which express some Muc1, but overall had a thinner glycocalyx than 1E7 and SKOV3(Singh et al., 2007). Against these targets, CAR-NK cells were more effective at killing than StcE-NK, although StcE-NK did kill a significantly higher percentage of SKBR3 cells compared to unmodified NK-92. Equipping the NK cells with both StcE and the CAR resulted in a modest enhancement of cytotoxicity compared to the StcE-NK and CAR-NK cells (Figure 5G). We also evaluated ovarian cancer OVCAR3 cells, which express Muc1 and the very large cell-surface mucin, Muc16(Coelho et al., 2018; Giannakouros et al., 2015) (Figure 5H). We could not accurately measure the glycocalyx thickness of OVCAR3 cells due to the poor adhesion of these cells to the SAIM substrate, which was possibly due to a thick mucin barrier on these cells. Mirroring the results for SKOV3, StcE-NK cells were more effective at killing these targets than CAR-NK. Collectively, these results indicate that immune cell engineering can help overcome the resistance to cellular attack posed by the mucin barrier.

## DISCUSSION

In summary, we have demonstrated that NK cell-mediated cytotoxicity shows a strong inverse relationship with the material thickness of the glycocalyx across widely varying states of glycosylation (Figure 4B). By combining gene editing techniques and the Ring-SAIM approach for nanoscale optical characterization of the glycocalyx, we demonstrated how specific molecular attributes of mucins, including their glycosylation and backbone length, contribute to the expansion and material thickness of the glycocalyx and, thus, the barrier to immune cell attack. We observed that equipping NK cells with a CAR or surface displayed StcE improved cytolytic activity against target cells. Whether CAR-NK or StcE is more effective may depend on the thickness of the mucin barrier, which is determined by the type and density of mucins on the cell surface.

In malignant conditions, differential glycosylation frequently results in the generation of truncated O-glycan structures, such as the Tn and sialyl-Tn antigens(Burchell et al., 2018). Our results reveal that mucins with truncated O-glycans can still resist close contact with an apposed surface when expressed at high levels. In general, increased mucin surface density can compensate for O-glycan truncation or loss of sialylation to maintain the structural integrity and thickness of the glycocalyx. Thus, the mucin barrier that is characterized here may be broadly relevant to cancer types that overexpress mucins, even across the highly heterogeneous patterns of glycosylation that are observed among specific tumor types and individual patients.

Our work has led to the development of quantitative relationships that describe the effect of mucin chain length and surface density on glycocalyx thickness. From the perspective of personalized medicine, mucin transcript levels, allele lengths, and surface levels are accessible quantities(Ashley, 2016; Tyanova et al., 2016). Combined with the simple scaling relations developed here, such metrics might inform the likelihood of a mucin barrier and, thus, patient recalcitrance to immune therapies. Few transferable standards exist for benchmarking and reproducibly categorizing mucin surface levels in clinical or research samples across labs. As one possible standard, the 1E7 cell line generated in this work allows titration of Muc1 levels with high reliability and reproducibility.

The studies here considered only a single physical metric of the glycocalyx barrier: its physical thickness. However, both the thickness and compressibility of the glycocalyx layer have long been predicted to resist the specific interactions of cells(Bell et al., 1984). Theoretical studies have shown that the deformation of the glycocalyx under compressive stress could vary considerably and in highly nonlinear ways depending on whether the mucin constituents have a flexible versus more rigid, rod-like structure(Gandhi et al., 2019). Muc1, which has only 5 consensus glycosylation sites per tandem repeat of 20 amino acids, behaves as a semi-flexible polymer(Shurer et al., 2019). Other cancer-associated mucins, including Podocalyxin, Muc4, and Muc21, have much higher glycosylation site density and may adopt more rigid structures compared to Muc1 when fully glycosylated(Nason et al., 2021). The potential importance of the mucin barrier in cancer immune evasion warrants a more complete physical characterization and understanding of the cell-surface mucinome.

Cancer immunotherapy and adoptive cell therapy have progressed rapidly, but clinical success has only been achieved in a small subset of cancers to-date(Waldman et al., 2020). Identifying and interfering with the mechanisms of cancer resistance to immunotherapy is an essential step towards continued progress. Strategies for disrupting the immunomodulatory glycocalyx have been proposed, including editing the glycocalyx structure with antibody-enzyme conjugates(Xiao et al., 2016). Here, we show that glycocalyx editing enzymes can be directly tethered to the immune cell surface to provide a performance benefit. The StcE-mucinase has a X409 lectin domain that can be used for linkage to the NK surface and possibly other types of clinically important effector cells. Other strategies for potentially more stable mucinase surface display include genetic encoding with a fused membrane anchor. Such strategies might be essential for in vivo translation of the findings described here. In view of the understanding of the glycocalyx as a material barrier against immune cell attack, strategies that can disrupt the glycocalyx or interfere with its synthesis are promising candidates for immunotherapy.

## MATERALS AND METHODS

### Cell lines

MCF10A cells were cultured in DMEM/F12 media (Thermo Fisher Scientific) supplemented with 5% horse serum (Thermo Fisher Scientific), 20 ng/mL EGF (Pepro Tech), 10 mg/ml insulin (Sigma), 500 ng/mL hydrocortisone (Sigma), 100 ng/mL cholera toxin (Sigma) and 1x penicillin/streptomycin (Thermo Fisher Scientific) at 37°C in 5% CO_2_. HEK293T cells was cultured in DMEM high glucose media (Thermo Fisher Scientific) supplemented with 10% fetal bovine serum (Thermo Fisher Scientific) and 1x penicillin/streptomycin (Thermo Fisher Scientific) at 37°C in 5% CO_2_. ZR-75-1 cells and T47D cells were cultured in RPMI media (Thermo Fisher Scientific) supplemented with 10% fetal bovine serum (Thermo Fisher Scientific) and 1x penicillin/streptomycin (Thermo Fisher Scientific) at 37°C in 5% CO_2_. SKOV-3 and SK-BR3 cells were cultured in McCoy’s 5a Medium Modified (Thermo Fisher Scientific) supplemented with 10% fetal bovine serum and 1x penicillin/streptomycin. NK-92 cells (ATCC CRL-2407) were cultured in ɑ-MEM without ribonucleosides media (Thermo Fisher Scientific) supplemented with 12.5% fetal bovine serum (Thermo Fisher Scientific), 12.5% horse serum (Thermo Fisher Scientific), 0.2 mM Myo-inositol (Sigma Aldrich), 0.1 mM 2-mercaptoethanol (Thermo Fisher Scientific), 0.02 mM folic acid (Millipore Sigma), 100 U/ml recombinant human IL-2 (Pepro Tech), and 1x penicillin/streptomycin (Thermo Fisher Scientific) at 37°C in 5% CO_2._

### Antibodies and Reagents

The following antibodies were used: anti-Human Muc1 Clone HMPV (555925, BD Biosciences), anti-Human Muc1 Janelia Fluor 549 (NBP-2-47883JF549, Novus Biologicals), Recombinant anti-GNE antibody (ab189927, Abcam), anti-GCNT1 antibody (ab102665, Abcam), anti-Galectin-3 N-20 (19280, Santa Cruz), anti-Galectin-1 (ab108389, Abcam), mouse anti-β-Actin Clone C4 (47778, Santa Cruz), anti-ErbB2/HER2 Clone 3B5 (ab16901, Abcam), APC conjugated anti-hErbB2/HER2 (FAB1129A, R&D systems), APC conjugated anti-Human perforin (Clone dG9, 30811, BioLegend), anti-ErbB2/HER2 Clone 3B5 (ab16901, Abcam), Anti-6X His-tag antibody (ab9108, Abcam). The following secondary antibodies were used: goat anti-mouse IgG DyLight 800 4x PEG conjugate (SA535521, Invitrogen), goat anti-mouse IgG DyLight 680 conjugate (35518, Invitrogen), goat anti-rabbit IgG Alexa Fluor 647 conjugate (A21245, Thermo Fisher Scientific), goat anti-mouse Alexa Fluor 647 conjugate (A28181, Invitrogen), and rabbit anti-goat IgG Alexa Fluor 647 conjugate (A21446, Invitrogen). Lectins used were: CF640R-conjugated PNA (#29063, Biotium), biotin-conjugated MAL-II (B-1265-1, Vector Laboratories). Biotinylated lectins were detected using NeutrAvidin Protein (DyLight 800 conjugated, 22853; DyLight 650 conjugated, 84607; both from Thermo Fisher Scientific). For tetracycline-inducible systems, doxycycline was used for induction (204734, Santa Cruz). For western blot, RIPA lysis buffer (89900, Thermo Fisher Scientific) and Halt Protease and Phosphatase Inhibitor Cocktail (78446, Thermo Fisher Scientific) were used. TD139 (28400, Cayman Chemical) was used to inhibit galectin-3 binding to Muc1. For digesting N-glycan, PNGase F (P0704S, New England Biolabs) was used. For detecting dead cell population by NK-mediated killing assay, propidium iodide (P4170, Sigma) and Annexin V conjugated CF647 (#29003, Biotium) were used. For SAIM measurement, MemGlow dye (MemGlow 560: MG02-2; MemGlow 590: MG03-10; MemGlow 640: MG04-02, Cytoskeleton) or CellBrite Steady 550 (#30107-T, biotium) were used to label cell membrane.

### Cloning and Constructs

cDNAs for mOxGFP-tagged Muc1 were previously cloned into a PiggyBac expression system containing a tetracycline-inducible promoter (P*_CMV-min_*-*tetO_7_*) and a separate puromycin selectin cassette (pPB tetOn PuroR)(Pan et al., 2019; Shurer et al., 2019). In this work, Muc1 cDNAs with approximately 42 native tandem repeats (TRs; Muc1 mOxGFP ΔCT) or either 0, 10, 21, and 42 perfect TRs of PDTRPAPGSTAPPAHGVTSA (0TR, 10TR, 21TR, 42TR) were used. Lentiviral plasmids for constitutive expression of human *GCNT1*, *LGALS3* (galectin-3), *HER2*, and an anti-HER2 chimeric antigen receptor (CAR) were prepared by inserting the cDNAs in place of GFP in pLenti CMV GFP Hygro (Addgene #17446), using the BamHI and BsrGI restriction sites. The unmodified human cDNA for *GCNT1* was generated by custom gene synthesis (General Biosystems). The human *HER2* cDNA was amplified by PCR from the plasmid, *HER2* WT (Addgene #16257), with primers that added 5’ BamHI and 3’ BsrGI restriction sites: 5’-TAATCTAGACCCAAGCTTGGGGCAG -3’ and 5’-GCTGTCGACGAGCTCGGTACCAAGCTT-3’. cDNA for human *LGALS3* was amplified by PCR from a pET21a expression vector (gift of C. Bertozzi) with primers that added 5’ BamHI and 3’ BsrGI restriction sites: 5’-TAAGCGGATCCGAATTCTATGGCAGACAATT-3’ and 5’-TGCTTTGTACATTATATCATGGTATATGAAGC-3’. cDNA for a previously reported HER2-specific CAR(Schönfeld et al., 2015) – including a humanized FRP5 scFV, CD8a hinge, CD28 transmembrane domain, CD28 signaling domain, and CD3ζ signaling domain – was synthesized with a 2A peptide/mTagBFP2 expression reporter by custom gene synthesis (Twist Bioscience) (See Supplemental Table 1 for the complete sequence). A pLentiCRISPR V2 plasmid for knockout of GNE was created by inserting the oligos, 5’-CACCGACATCAAGTTCAAAGAACTC -3’ and 5’-AAACGAGTTCTTTGAACTTGATGTC-3’, into the BsmBI restriction site following standard protocols.(Sanjana et al., 2014) Recombinant GFP nanobody cDNA (PDB #3OGO_E) was generated by custom gene synthesis (IDT) and cloned into the BamHI and HindIII restriction sites of pTP1112 (Addgene #104158).

### Generated Cell Lines

MCF10A cells stably expressing the rtTA-M2 tetracycline transactivator were prepared by lentiviral transduction using the pLV rtTA-NeoR plasmid as previously described.(Paszek et al., 2012) For preparation of mucin-expressing cell lines, MCF10A rtTA-M2 cells were co-transfected with the PiggyBac hyperactive transposase and PiggyBac plasmids containing ITR-flanked expression cassettes (i.e. pPB tetOn PuroR plasmids) using the NEPA21 Electroporator (NEPAGENE) and subsequently selected with 1 µg/mL puromycin. The stable MCF10A line expressing native Muc1 mOxGFP ΔCT was clonally expanded by limiting dilution of single cells in 96-well plates. An individual clone, 1E7, was isolated that exhibited low leaky expression of Muc1 mOxGFP ΔCT and titratable Muc1 surface levels with doxycycline induction.

Progeny of the 1E7 clone overexpressing *GCNT1*, *LGALS3*, and *HER2* were prepared by transduction with lentiviral particles generated using the respective pLenti CMV Hygro plasmids. Cell selection was with 200 µg/mL hygromycin. Control MCF10A rtTA-M2 cells overexpressing *HER2* were similarly prepared. Knockout (KO) of *GNE* in the 1E7 clone was achieved by lentiviral transduction using the pLentiCRISPR V2 system and selection with 200 µg/mL hygromycin. The selected cells were clonally expanded and individual clones were screened for double knockout of *GNE* by Sanger sequencing of genomic DNA that was isolated using the Quick-DNA miniprep kit (Zymo, D3024).

KO of *C1GALT1* in the 1E7 cells was achieved using the Alt-R CRISPR/Cas9 system (IDT) with homology-directed repair (HDR) templates that introduced an in-frame stop codon and an EcoRI restriction site for screening. CRISPR RNA (crRNA) and HDR template for *C1GALT1* KO were: 5’-GTAAAGCAGGGCTACATGAG-3’ and 5’-CTTTGGGAGAAGATTTAAGCCTTATGTAAAGCAGGGCTACTAAGAATTCAGTGGAGGAGCAGGATATGTACTAAGCAAAGAAGCCTTGA-3’. CRISPR/Cas9 editing was conducted according to manufacturer’s protocol for the Alt-R system. Briefly, 1E7 cells were transfected with precomplexed crRNA and ATTO550-tracrRNA, along with the HDR template, using electroporation. After 48 hours, transfected 1E7 cells were clonally expanded by limiting dilution in 96-well plates. To screen for successful knockouts, genomic regions encompassing the targeted CRISPR/Cas9 cut sites were amplified by PCR and analyzed by EcoRI restriction digest and Sanger sequencing. PCR primers for *C1GALT* were 5’-GGAGGATAATAGTTGTAATTCCAGTACCAAAAC-3’ and 5’-TCAAAACCTAGAGAAAAAGGCCAAACAC-3’. PCR primers for GNE were 5’-AGTGGTTAAGGACTTGAAACTG-3’ and 5’-TCTACTAAGCGGCATCATTG-3’. Sequencing primer for *C1GALT1* was 5’-ACACGTCAAAGCTACTTGG-3’ and sequencing primer for *GNE* was 5’-AGTGGTTAAGGACTTGAAACTG-3’. A NK-92 line expressing the HER2-CAR was prepared by transduction with lentiviral particles generated using the pLenti CMV Hygro plasmid. Cell selection was with 200 µg/mL hygromycin. Selected cells were sorted for high mTagBFP2 signal.

### Western Blot Analysis

Cells were plated and induced with 1 μg/mL doxycycline for 24 hours before lysis with RIPA buffer. Lysates were separated on NuPAGE 3-8% Tris-Acetate gels or NuPAGE 4-12% Bis Tris gels and transferred to low-fluorescence PVDF membranes (Milipore Sigma, IPFL07810). Primary antibodies were diluted at 1:500 and fluorophore-conjugated or biotinylated lectins were diluted to 2 μg/mL in 5% BSA TBST and incubated overnight at 4°C. Secondary antibodies and Neutravidin-DyLight 800 were diluted at 1:1000 or 1 μg/mL in 5% BSA TBST and incubated for 1 hour at room temperature. Blots were imaged on a ChemiDoc MP Imaging System (Bio-Rad)

### Flow Cytometric Analysis

Cells were plated, grown for 24 hours, and then induced with various concentrations of doxycycline for 24 hours. Adherent cells were detached by incubating with Accutase (Innovative Cell Technologies) at 37°C for 10 minutes. GFP nanobody conjugated Alexa Fluor 647, Neutravidin conjugated DyLight 650, CF640R PNA, Alexa Fluor 647 HPA, and biotin MAL-II were diluted 1:200 in 0.5% BSA PBS and incubated with cells at 4°C for 30 minutes for each stain. For analysis of Muc1 cell surface expression level, anti-Human Muc1 Clone HMPV was diluted 1:200 in 0.5% BSA PBS and incubated with cells at 4°C for 1 hour. Secondary labelling was with Alexa Fluor 647 conjugated goat anti-mouse, diluted 1:1000 in 0.5% BSA PBS and incubated with cells at 4°C for 1 hour. For analysis of StcE mucinase binding to NK-92 cells, NK cells were incubated with StcE at 37°C for 1 hour. Cells were thoroughly washed to remove StcE and coculture with target cells. Anti-6X His-tag antibody was diluted 1:100 in 0.5% BSA PBS and incubated with cells at 4°C for 1 hour. Secondary labelling was with Alexa Fluor 647 goat anti-rabbit IgG, diluted 1:500 in 0.5% BSA PBS and incubated with cells at 4°C for 1 hour. To analyze fluorescent labelled cells, Attune NxT Flow cytometry (Thermo Fisher) was used.

### Immunofluorescence

MCF10A cells were plated and induced with 1, 100, and 1,000 ng/mL doxycycline for 24 hours. For endogenous galectins staining of cells, galectin antibodies were diluted 1:50 with 0.5% in 0.5% BSA PBS and incubated on 1E7 clone with 1,000 ng/mL of doxycycline at 4°C for 1 hour after being fixed with 4% paraformaldehyde and permeabilized with 0.1% Triton X-100. For cell-surface galectin-3, TD139 (28400, Cayman Chemical) was diluted to 1, 10, 50 µM in MCF10A media and incubated on 1E7 clone with 1,000 ng/mL of doxycycline at 37°C for 2 hours. TD139-treated cells were further incubated with anti-galectin-3 antibody at 4°C for 30 minutes. For GCNT1 imaging, 1E7 clone cells were fixed with 4% paraformaldehyde and permeabilized with 0.1% Triton X-100 after induction of 1,000 ng/mL doxycycline. GCNT1 antibody was diluted 1:50 in 0.5% BSA PBS and incubated on samples at 4°C for 1 hour. For NK-92 cell imaging, NK-92 cells were incubated with 100 nM StcE in NK cell culture media for 1 hour at 37°C and washed thoroughly with phenol red-free DMEM/F12 supplemented with 5% horse serum, 20 ng/mL EGF, 10 mg/ml insulin, 500 ng/mL hydrocortisone, 100 ng/mL cholera toxin and 1x penicillin/streptomycin. 1E7 cells were plated to glass bottom dishes for 1 days and induced with 1,000 ng/mL doxycycline for 24 hours. StcE-treated NK cells were plated on 1E7 cells at 37°C for 4 hours. For Hyaluronic acid imaging, cells were incubated with 2.5 µg/mL Hyaluronic Acid Binding Protein Biotinylated (385911, Millipore Sigma) at 4°C for 1 hour. For other Muc1-GFP clones imaging, two other clones (2E4 and 2G9) were plated onto glass-bottom dishes and induced with 1, 100, 1,000 ng/mL of doxycycline (sc-204734, Santa Cruz) for 24 hours. Secondary antibodies were diluted 1:500 in 0.5% BSA PBS and incubated with cells at 4°C for 1 hour. Samples were imaged on a LSM 800 confocal microscope using 10x (NA: 0.3 Air), 20x (NA: 0.8 Air), and 63x (NA: 1.4 Water) objectives (Zeiss).

### Preparation of recombinant human galectins

Human *LGALS1* and *LGALS3* constructs in pET21 were recombinantly expressed in *E. coli* strain NiCo21 (DE3) (New England Biolabs). Transformed bacteria were grown in LB at 37°C until an OD600 of 0.6 – 0.8 was reached. Expression was induced with 0.3 mM IPTG, and protein was produced overnight at 20°C. Cells were harvested by centrifugation at 3,000 g for 20 minutes and lysed by a pressurized homogenizer with cOmplete protease inhibitor Cocktail (Roche) and 1 mg/mL lysozyme (Sigma). The lysate was centrifuged at 20,000 rpm for 45 minutes at 4°C, and the supernatant was incubated with β-lactosyl Sepharose resin for 1 hour at 4°C before loading into a gravity column. The protein was eluted with 0.1 M β-lactose (Santa Cruz Biotech) and 8 mM DTT (Sigma). The partially purified protein was polished on a HiPrep 16/60 Sephacryl S-100 HR (GE Healthcare Life Sciences) column equilibrated with 0.1 M β-lactose and 8 mM DTT. The final protein was then concentrated by using Amicon Ultra (3kD MWCO for Galectin-1; 10kD MWCO for Galectin-3) filters (Millipore Sigma). Conjugation of β-lactose to Sepharose 6B (Sigma) to synthesize the β-lactosyl Sepharose was as previously described(Pace et al., 2003).

### StcE mucinase Expression and Purification

The cDNA for StcE-Δ35(Yu et al., 2012) was synthesized by custom gene synthesis (Twist Bioscience) and inserted into the pET28b expression vector. StcE E447D was generated using Q5 Site-Directed Mutagenesis Kit (New England Biolabs) with primers 5’-TCAGTCATGACGTTGGTCATAATTATG-3’ and 5’-ACTCATTCCCCAATGTGG-3’. StcE-ΔX409 was also generated using the Q5 Site-Directed Mutagenesis Kit with primers 5’-TAACTCGAGCACCACCAC-3’ and 5’-ATTTACAGTATAGGTAAGTCCTTC-3’. Cells were harvested by centrifugation at 3,000 g for 20 minutes, resuspended in lysis buffer (20 mM HEPES, 500 mM NaCl and 10 mM imidazole, pH 7.5) with cOmplete protease inhibitor Cocktail (Roche), and lysed by a pressurized homogenizer. StcE was purified by immobilized metal affinity chromatography (IMAC) on a GE ÄKTA Avant FPLC system. The lysate was loaded onto a HisTrap HP column (GE Healthcare Life Sciences), washed with 20 column volumes of wash buffer (20 mM HEPES, 500 mM NaCl and 20 mM imidazole, pH 7.5.), and eluted with a linear gradient of 20 mM to 250 mM imidazole in buffer (20 mM HEPES and 500 mM NaCl, pH 7.5.). The elution fractions containing target protein were collected and polished on a HiPrep 26/60 Sephacryl S-200 HR (GE Healthcare Life Sciences) column equilibrated with storage buffer (20 mM HEPES and 150 mM NaCl, pH 7.5). The final protein was then concentrated by using Amicon Ultra 30 kDa MWCO filters (Millipore Sigma)(Malaker et al., 2019).

### GFP Nanobody Preparation and Conjugation

Recombinant GFP nanobody (PDB #3OGO_E) was prepared in chemically competent NiCo21 (DE3) E. coli (NEB) using the pTP1112 plasmid. Transformed bacteria were grown in LB at 37°C until an OD600 of 0.6 was reached. The cultures were induced with 0.5 mM IPTG overnight at 24°C, harvested, resuspended in B-PER (Thermo Fisher Scientific) and vortexed for cell lysis. The lysates were cleared by centrifugation at 10,000 g for 20 minutes at 4°C. The His-tagged nanobody was purified by IMAC according to standard protocols. Briefly, supernatant diluted into 1x Ni-NTA binding buffer was bound to equilibrated Ni-NTA resin (Qiagen, 30210) for 20 minutes at 4°C, with end-over-end mixing. The resin was added to a spin column, washed thoroughly, and incubated with the Ni-NTA elution buffer for 20 minutes at 4°C, mixing end-over-end. Eluted protein was exchanged into storage buffer (pH 7.4 PBS) using Zeba 7K MWCO desalting columns or by overnight dialysis with 10K MWCO Snakeskin dialysis tubing. Eluted proteins were then sterile filtered and snap-frozen for long-term storage at -80°C. Nanobody-containing constructs were mixed with 0.1% w/v sodium azide prior to snap-freezing. For fluorescent GFP nanobody, purified GFP nanobody was labeled with Alexa Fluor 647 NHS Ester (Thermo Fisher Scientific) per the manufacturer’s protocol.

### NK Cell Cytotoxicity Assay

Target MCF10A cells were fluorescently labeled by incubation with 1 μM CellTracker Green CMFDA Dye (Invitrogen) in MCF10A growth media for 10 minutes. 3×10^4^ labeled target cells were suspended with varying ratios of NK-92 effector cells in 200 µL of growth media of target cell line in absent of IL-2 and cocultured in an ultra-low attachment U-bottom 96-well plate (Corning) for 4 hours at 37°C in 5% CO_2_. For all StcE-treated NK cell experiment, unless otherwise indicated, NK cells were incubated with 100 nM StcE at 37°C for 1 hour. Cells were thoroughly washed to remove StcE and coculture with target cells. After 4 hours, propidium iodide (PI; 20 μg/mL, Sigma) was added to each well for 10 minutes and washed with 0.5% BSA PBS. For different Muc1 surface level on ZR-75-1, anti-Human Muc1 Janelia Fluor 549 was diluted 1:200 in 0.5% BSA PBS and incubated with ZR-75-1 cells at 4°C for 1 hour and sorted the cells with defined standardized gates for low, moderate, and high Muc1 expression by FACS (Sony MA900 Cell Sorter). Sorted target cells were cocultured with varying ratios of NK-92 effector cells for 4 hours. Annexin V conjugated with CF647 (0.5 μg/mL, Biotium) was diluted in HEPES-buffered saline containing 2.5 mM calcium chloride and added to each well for 15 minutes. NK cell cytotoxicity was evaluated by flow cytometry as previously described(Bryceson, 2010; Hudak et al., 2014a). At least 1×10^4^ target cells were analyzed after electronic gating on CellTracker Green. Percent cytotoxicity was calculated as 100 × (experimental % dead − spontaneous % dead)/(100 − spontaneous % dead), where experimental % dead was the percentage of PI positive target cells in NK cocultures and spontaneous % dead was the percentage of PI positive control target cells cultured in the absence of effector cells.

### Fluorescence recovery after photobleaching

FRAP experiments were performed by using a Zeiss i880 confocal microscope with 63x N.A. 1.4 Oil immersion lens. Five pre-bleached images were firstly taken and photobleaching was performed with 488 nm laser for about 10 seconds to bleach a target area of 2.55 µm x 2.55 µm. 1E7 clone (n=11) and 1E7 GCNT1 overexpression (n=10) were used to measure mobile fraction of Muc1-GFP. 10 µM recombinant human galectin-1 was incubated with 1E7 GCNT1 cells for 10 minutes at 37°C (n=9). Intensity profiles were corrected by normalizing to the time-course decay of GFP signal in non-bleached area by using MATLAB code from Advanced Imaging Center (https://www.mathworks.com/matlabcentral/fileexchange/47327-frap-zip)

### Scanning Angle Interference Microscopy

#### Sample preparation

Silicon wafers with ∼1900 nm thermal oxide (Addison Engineering) were diced into 7 mm by 7 mm chips, and the oxide layer thickness for each chip was measured with a FilMetrics F50-EXR. Prior to use, the chips were then cleaned with 1:1 methanol and hydrochloric acid for 15 minutes, followed by 1 minutes in a plasma cleaner (Harrick Plasma, PDC-001). The chips were incubated with 4% (v/v) (3-mercaptopropyl)trimethoxysilane in absolute ethanol for 30 minutes at room temperature. After washing with absolute ethanol, the chips were subsequently incubated with 4 mM 4-maleimidobutyric acid N-hydroxysuccinimide ester in absolute ethanol for 30 minutes and rinsed with PBS. 50 µg/mL human plasma fibronectin in PBS was incubated with the functionalized chips overnight at 4°C for conjugation. Cells were seeded onto the chips at 2 × 10^4^ cm^−2^ in full culture medium with varying doxycycline concentrations and incubated for 24 hours. Cells expressing different length of Muc1 were labelled with GFP nanobody conjugated to Alexa Fluor 647 in MCF10A growth media for 10 minutes at 37°C. Cells were labelled with MemGlow dye or CellBrite Steady 550 in serum-free, phenol red-free DMEM/F12 for 10 minutes at 37°C. Sample were then rinsed 3x in PBS, inverted into a 35 mm glass-bottom imaging dish containing imaging buffer (MemGlow dye or CellBrite dye: serum-free, phenol red-free DMEM/F12, Alexa Fluor 647-conguated GFP nanobody: phenol red-free DMEM/F12 supplemented with 5% horse serum, 20 ng/mL EGF, 10 mg/ml insulin, 500 ng/mL hydrocortisone, 100 ng/mL cholera toxin and 1x penicillin/streptomycin) and imaged at 37°C. For NK-92 cell, the cleaned chips were incubated with 2.5% (v/v) (3-mercaptopropyl)trimethoxysilane, 4.5% (v/v) deionized water and 0.9% (v/v) acetic acid) in methanol overnight at 4°C. After washing with PBS, the chips were reacted with 0.1 mg/mL of maleimide-activated neutravidin protein (31007, Thermo Scientific) with 50 μg/mL fibronectin for 1 hour at room temperature and rinsed with PBS. A biotin-conjugated MAL-II were diluted 1:200 in PBS and incubated with the chips for 1 hour at room temperature. NK-92 cells were seeded onto the chips at 2 × 10^4^ cm^−2^ in full culture medium and incubated for 1 hour(Kim et al., 2018).

#### Optical setup

Scanning angle interference microscopy (SAIM) was conducted on a custom circle-scanning microscope.(Colville et al., 2019) The core of the setup was an inverted fluorescence microscope (Ti-E, Nikon). The excitation lasers (488 nm, Coherent; 560 nm, MPB Communications Inc.; and 642 nm, MPB Communications Inc.) were combined into a colinear beam by a series of dichroic mirrors (Chroma). The combined output beam was attenuated and shuttered by an AOTF (AA Opto-Electronic). The beam was directed onto a pair of galvanometer scanning mirrors (Cambridge Technology). The image of the laser on the scanning mirrors was magnified and relayed to the sample by two 4 f lens systems, a beam expanding telescope and a scan lens / objective lens combination. The beam expander was formed by f = 30 mm and f = 300 mm achromatic lenses with a m = 1 zero-order vortex half-wave plate positioned between them and positioned 2 f from the 300 mm achromatic scan lens (Thorlabs). SAIM experiments were performed with a 60x N.A. 1.27 water immersion objective (Nikon). Fluorescence emission was collected with a quad band filter cube and single band filters (TRF89901-EMv2, Chroma) mounted in a motorized filter wheel (Sutter). Images were acquired with a Zyla 4.2 sCMOS (Andor) camera or an iXon 888 EMCCD (Andor) using the microscope’s 1.5x magnifier for a total magnification of 90×. The open-source software Micro-Manager was used for camera and filter wheel control and image acquisition. The circle-scanning galvometers were operated in an autonomous fashion using a custom-designed 16-bit PIC microcontroller, which has been described previously(Colville et al., 2019).

#### Image Acquisition and Analysis

For SAIM, 32 images were acquired at varying incidence angles for the circle-scanned excitation beam. During a typical image acquisition sequence, changes in the scanned incidence angle were triggered by a TTL signal from the camera to the microcontroller. The incidence angles were evenly spaced from 5 to 43.75 degrees with respect to the wafer normal in the imaging media. To obtain the reconstructed height topography of the samples, the raw image intensities, I_j_, at each incidence angle, θ_j_, were fit pixelwise by nonlinear least-squares to an optical model:

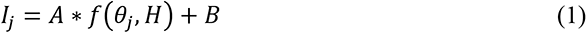

where H is the unknown sample height and A and B are additional fit parameters. The vortex half-wave plate in the optical setup maintained the s-polarization of the circle-scanned excitation laser. For s-polarized monochromatic excitation of wavelength, λ, the probability of excitation, *f*(*θ_j_*, *H*), for the system is given by:

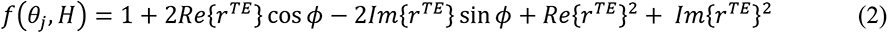

where the phase shift, ϕ, and the reflection coefficient for the transverse electric wave, *r^TE^*, are given by:

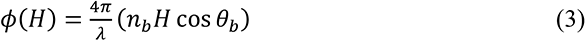

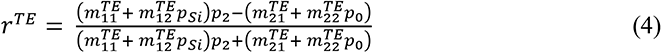

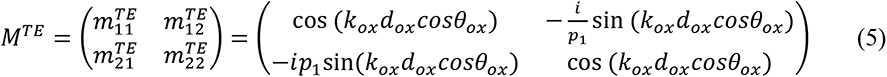

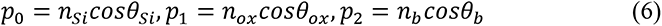

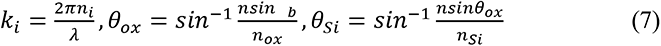

where *k_i_* is the wavenumber in material i; *n_Si_*, *n_ox_* and *n_b_* are the refractive index of the silicon, silicon oxide and sample, respectively; *θ_Si_*, *θ_ox_* and *θ_b_* are the angles of incidence in the silicon, silicon oxide and sample, respectively; and *d_ox_* is the thickness of the silicon oxide layer. The angles of incidence in silicon oxide and silicon were calculated according to Snell’s Law. To quantify the glycocalyx thickness of cells, the average height above the silicon substrate was calculated for a 100 x 100 pixel subregion in each cell. The glycocalyx thickness was reported as the height of the localized GFP nanobody, MemGlow dye or CellBrite dye signal minus the height of the fluorescently labeled fibronectin layer on the silicon substrate. Packages for fitting SAIM image sequences with the above model have been implemented in C++ and Julia and are available through GitHub: https://github.com/mjc449/SAIMscannerV3 and https://github.com/paszeklab/SAIMFitKit.jl

## ACKNOWLEDGMENT

This investigation was supported by National Institute of Health New Innovator DP2 GM229133 (M.J.P.), National Cancer Institute (NCI) U54 CA210184 (M.J.P. and C.F.), NCI R33-CA193043 (M.J.P and W.R.Z), National Institute of General Medical Sciences (NIGMS) R01 GM138692 (M.J.P), NIGMS R01 GM137314 (M.J.P and M.D.), and National Science Foundation 1752226 (M.J.P.). NIGMS R01 GM138692 supported the glycoengineered cell line development and glycocalyx material characterization. NCI U54 CA210184 supported the investigation of immune cell interactions with engineered and tumor cell lines. NCI R33-CA193043 supported the development of the SAIM microscope with additional support from the Kavli Institute at Cornell for Nanoscale Science. This work was performed in part at the Cornell NanoScale Facility, a member of the National Nanotechnology Coordinated Infrastructure (NNCI), which is supported by the National Science Foundation (Grant NNCI-2025233). Flow cytometry data was acquired through the Cornell University Biotechnology Resource Center (RRID:SCR_021740). FRAP imaging data was acquired through the Cornell Institute of Biotechnology’s Imaging Facility (RRID:SCR_021741), with NYSTEM (C029155) and NIH (S10OD018516) funding for the shared Zeiss LSM880 confocal/multiphoton microscope.

## AUTHOR CONTRIBUTIONS

All authors contributed to the design of experiments, interpretation of results, and preparation of the manuscript. M.J.C constructed the Ring-SAIM microscope. S.P. and M.J.C. conducted the Ring-SAIM measurements and analysis. S.P., C.R.S. and J.S. designed and constructed the mucin expression plasmids and other expression vectors. M.J.C. isolated and validated the 1E7 clone. S.P. and J.P. designed the CRISPR guide RNAs and HDR sequences. S.P. conducted all cytotoxicity assays. S.P. and M.J.P. wrote the manuscript with feedback from all authors.

## AUTHOR INFORMATION

Matthew J. Paszek

Address: 120 Olin Hall, 113 Ho Plaza, Cornell University, Ithaca NY 14853, USA

