## Supplementary Figures for "Mucins form a nanoscale material barrier against immune cell attack"

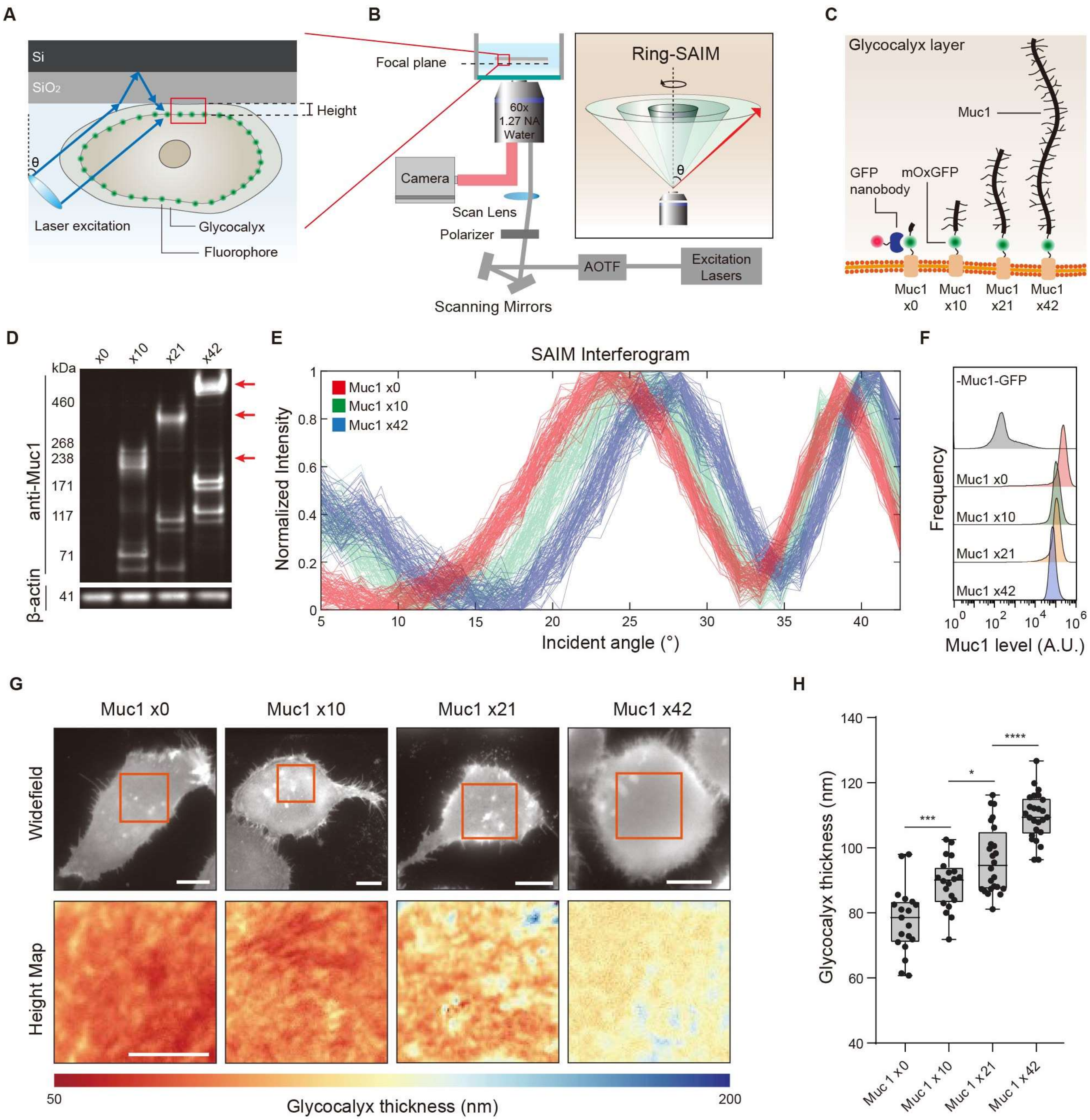


**Figure S1. Measurement of the glycocalyx material thickness with Ring Scanning Angle Interference Microscopy (Ring-SAIM).** (A) Schematic illustration of Ring-SAIM; interference of direct and reflected laser light off a silicon mirror create standing waves of excitation light that depend on the laser incidence angle, θ, and are used to probe the vertical position of fluorescence emitters. (B) Optical configuration for Ring-SAIM showing azimuthally scanned excitation light at varying θ; AOTF, Acousto-optic tunable filter; Polarizer, m = 1 vortex half-wave retarder. (C) Muc1-GFP constructs with biopolymer domains comprised of 0, 10, 21, and 42 tandem repeats (TRs); GFP-nanobody specifically labels the cell-surface fraction of Muc1-GFP. (D) Western blot analysis of Muc1-GFP constructs stably expressed in MCF10A epithelial cells; arrows indicate fully glycosylated Muc1 with 10, 21, or 42 TRs. (E) Examples of the pixelwise Ring-SAIM interferograms for Muc1-GFP constructs labelled with AF647-GFP nanobody. (F) Flow cytometry analysis of cell-surface Muc1-GFP probed with Alexa Fluor 647 (AF647) conjugated GFP nanobody. (G) Representative widefield image and glycocalyx thickness map of live epithelial cells expressing the indicated Muc1 constructs labeled with AF647 GFP nanobody (scale bars, 10 µm). (H) Quantification of mean glycocalyx thickness in cells expressing the Muc1 constructs. Results are the mean ± s.d. of at least 19 cells. Statistical significance is given by * P ≤0.05, *** P ≤ 0.001, **** P ≤ 0.0001. Results are a representative dataset from 3 independent replicates.

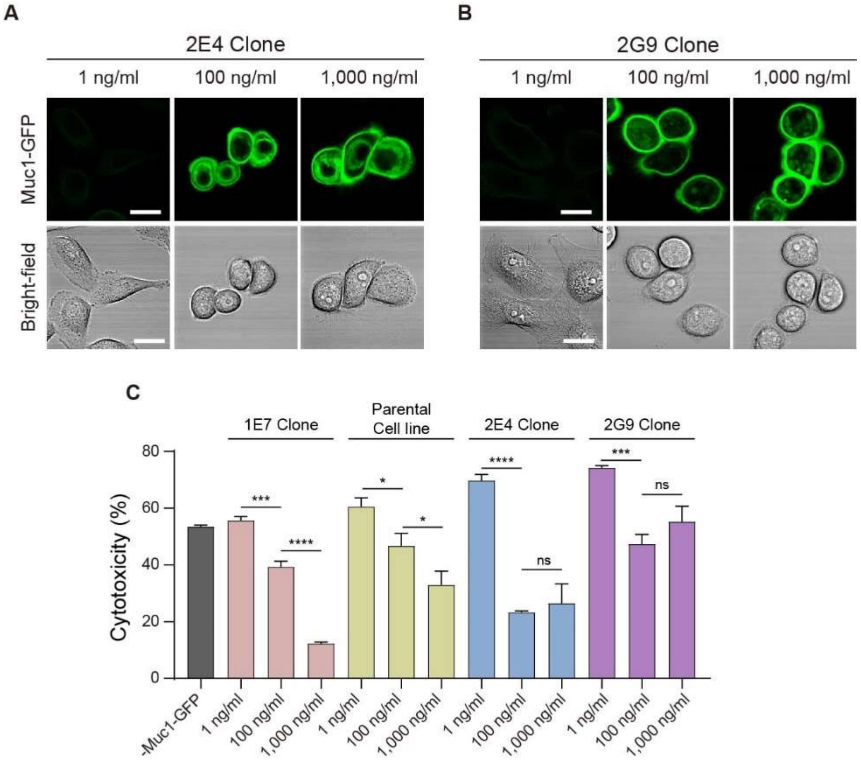


**Figure S2.** Analysis of NK-cell mediated killing in additional Muc1-GFP expressing clones. (A,B) Representative confocal and bright-field images of two clonally expanded MCF10A cell lines expressing Muc1-GFP with 42 tandem repeats under the control of a tetracycline inducible promoter; the doxycycline induction levels are indicated. (Scale bar, 10 µm). (C) Cytotoxicity of NK-92 cells against three clonally expanded cell populations and the parental polyclonal cell line. The doxycycline induction levels are indicated. NK cell to target cell ratio is 5:1.

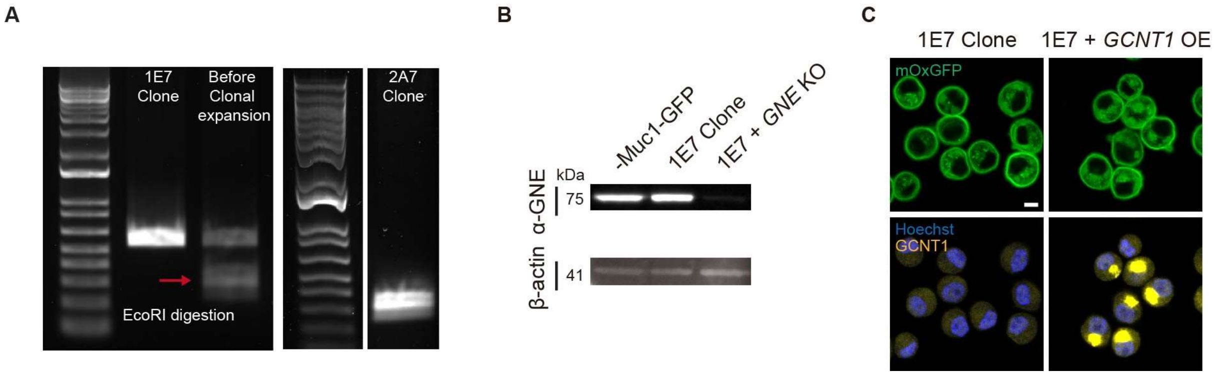


**Figure S3.** Validation of glycoengineered cell lines. (A) Agarose gel electrophoresis of PCR-amplified and EcoRI-digested segment *C1GALT1* showing successful homozygous knockout (KO) in 1E7 cells (See 2A7 sub-clone of 1E7). For the KO, a stop codon and an EcoRI restriction site for screening were inserted in the *C1GALT1* gene via CRISPR/Cas9 and homology directed repair. (B) Western blot analysis confirming UDP-N-acetylglucosamine 2-epimerase (*GNE*) KO in 1E7 cells. (C) Fluorescence images of GCNT1 in 1E7 clone and *GCNT1* overexpression in the 1E7 clone induced at 1,000 ng/ml of doxycycline (Scale bar, 10 µm).

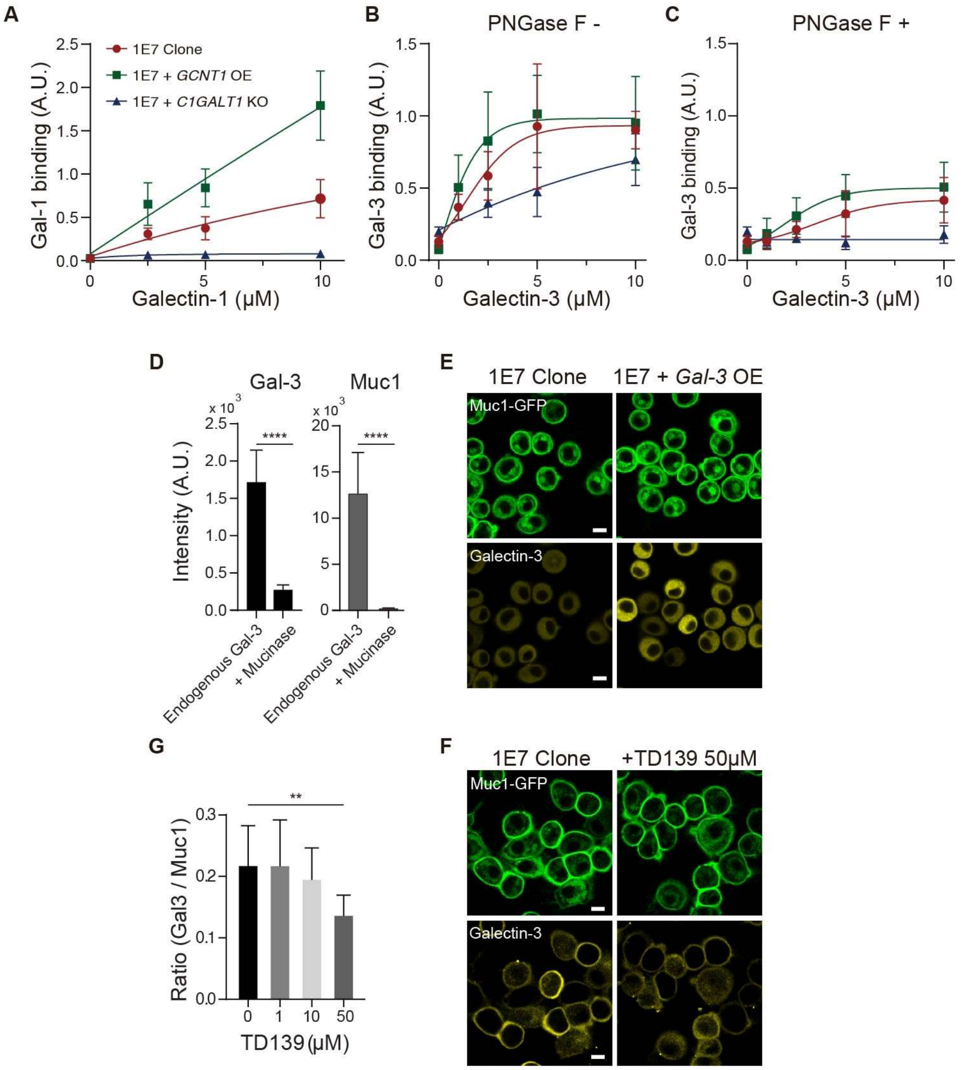


**Figure S4.** Confirmation of galectin-1 and galectin-3 interactions with mucin-type O-glycans (A) Quantification of recombinant galectin-1 (Gal-1) binding to Muc1-GFP expressing 1E7 cells and their glycoengineered progeny with knockout of C1GALT1 to ablate Core-I extension of O-glycans and overexpression (OE) of GCNT1 to increase Core-II extension; induction of Muc1-GFP is at 1,000 ng/ml of doxycycline; data normalized to the Muc1 signal intensity and presented as the mean and ± s.d. of 20 cells (B) Quantification of recombinant galectin-3 (Gal-3) binding to the 1E7 cells and their glycoengineered progeny; data normalized to the Muc1 signal intensity and presented as the mean and ± s.d. of 20 cells. (C) Same as in b with 5,000 U/mL PNGase F to selectively removes N-glycans prior to analysis. (D) Quantification of endogenous galectin-3 (left) and Muc1 (right) on cell surface before and after 200 nM of StcE mucinase treatment, confirming that galectin-3 binds specifically to Muc1. (E) Representative confocal images of permeabilized 1E7 clone and *Gal-3* overexpression in the 1E7 clone induced at 1,000 ng/ml of doxycycline (Scale bar, 10 μm). (F) Representative confocal images of endogenous galectin-3 on the cell membrane in presence and absence of TD139 (Scale bars, 10 μm). (G) Quantification of galectin-3 (Gal3) binding to 1E7 clone surface induced 1,000 ng/ml of dox with TD139; data normalized to the Muc1 signal intensity and presented as the mean and ± s.d. of 10 cells. Statistical significance is given by ** P ≤0.01, **** P ≤ 0.0001.

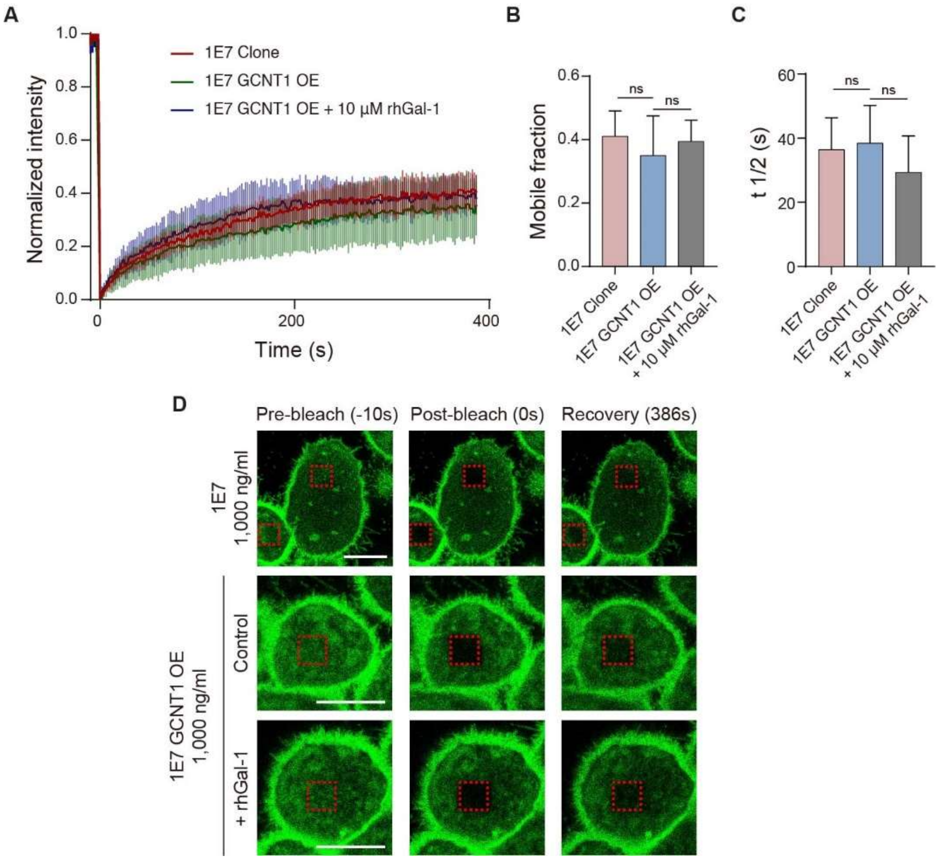


**Figure S5.** Galectin-1 does not significantly affect the mobility and mobile fraction of cell surface Muc1. (A) FRAP analysis was performed on 1E7 clone (red), 1E7 GCNT1 OE (green), 1E7 GCNT1 OE with 10 µM recombinant human galectin-1 (blue). n = 11, 10, 9 respectively (B,C) Quantitative analysis of mobile fraction (B) and half recovery time (C). (D) Representative confocal images of each condition at pre-bleach, post-bleach, and recovery. (Scale bar, 10 µm).

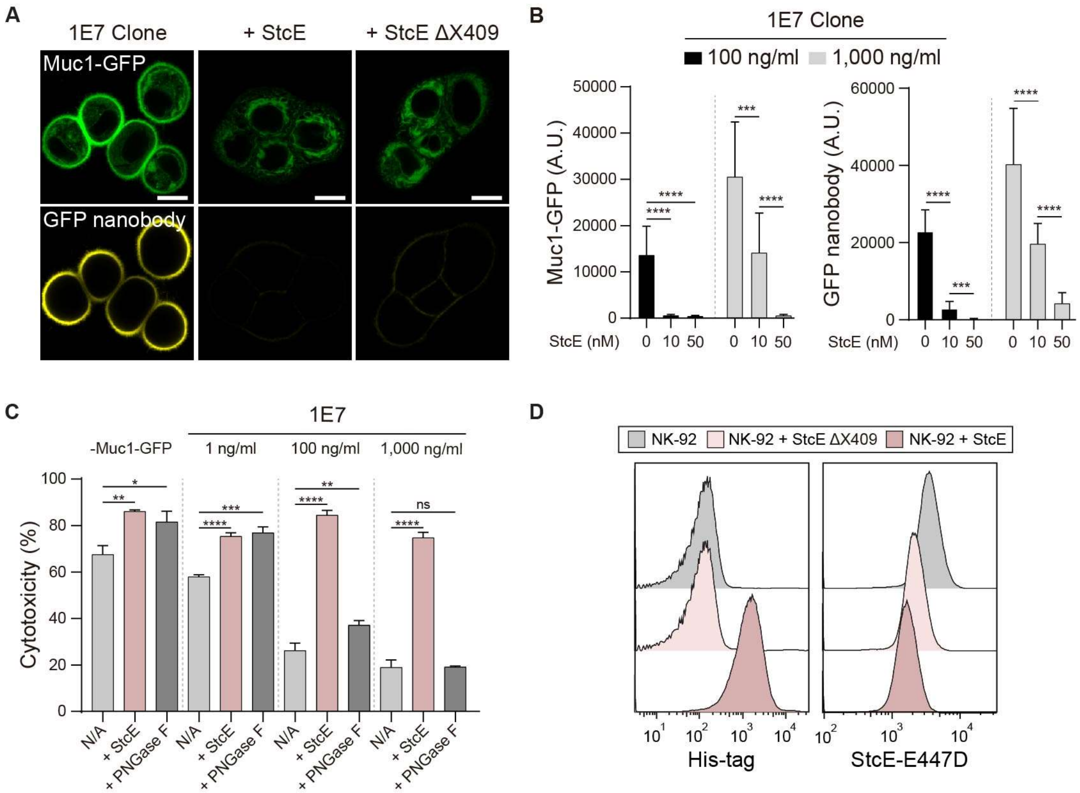


**Figure S6.** StcE mucinase effectively remove cell-surface mucin on 1E7 clone and NK cell. (A) Representative confocal images of Muc1-GFP expressing 1E7 cells with and without treatment with 100 nM StcE or StcE ΔX409; Alexa Fluor 647 conjugated to GFP nanobody is used to probe cell-surface Muc1-GFP constructs. (Scale bar, 10 μm). (B) Quantification of mean fluorescence intensity of Muc1-GFP and cell-surface Muc1-GFP probed with GFP nanobody in 1E7 cells treated with the indicated concentration of StcE mucinase; results are shown for induction of Muc1-GFP at 100 ng/ml and 1000 ng/ml doxycycline. (C) Cytotoxicity of NK-92 cells against 1E7 cells treated with control buffer, 100 nM StcE mucinase, or 5,000 U/mL PNGase F for 1 hour; Muc1-GFP induction is at the indicated doxycycline concentration (1,100, and 1,000 ng/ml); NK cell to target cell ratio is 5:1. (D) Flow cytometry analysis of mucin levels on NK-92 cells following incubation with control buffer or 100 nM recombinant StcE or StcE ΔX409; StcE surface level after incubation probed with anti His-tag antibody and cell-surface mucin level probed with catalytically dead StcE E447D conjugated to Cy5. Data presented as the mean and ± s.d. of three independent experiments. Statistical significance is given by **** P ≤ 0.0001.

**Table S1. Complete sequence for HER2-specific CAR**(Zhang et al., 2017)

HER2-specific CAR includes FRP5 scFv, CD8a Hinge, CD28 TM, and CD Zeta. mTagBFP2 sequence is followed by P2A sequence after CD3 Zeta.

| | |
|---|---|
| FRP5 scFV | CAGGTGCAGCTGCAGCAGAGCGGCCCTGAGCTGAAGAAGCCCGGCG AGACAGTCAAGATCAGCTGCAAGGCCAGCGGCTACCCCTTCACCAAC TACGGCATGAACTGGGTGAAACAGGCCCCAGGCCAGGGACTGAAGT GGATGGGCTGGATCAACACCAGCACCGGCGAGAGCACCTTCGCCGA CGACTTCAAGGGCAGATTCGACTTCAGCCTGGAAACCAGCGCCAACA CCGCCTACCTGCAGATCAACAACCTGAAGAGCGAGGACAGCGCCAC CTACTTTTGCGCCAGATGGGAGGTGTACCACGGCTACGTGCCCTACT GGGGCCAGGGCACCACCGTGACCGTGTCCAGCGGCGGAGGGGGCTC TGGCGGCGGAGGATCTGGGGGAGGGGGCAGCGACATCCAGCTGACC CAGAGCCACAAGTTTCTGAGCACCAGCGTGGGCGACCGGGTGTCCAT CACCTGCAAAGCCAGCCAGGACGTGTACAACGCCGTGGCCTGGTATC AGCAGAAGCCTGGCCAGAGCCCCAAGCTGCTGATCTACAGCGCCAGC AGCCGGTACACCGGCGTGCCCAGCAGGTTCACCGGCAGCGGCAGCG GCCCAGACTTCACCTTCACCATCAGCAGCGTGCAGGCCGAGGACCTG GCCGTGTACTTCTGCCAGCAGCACTTCCGGACCCCCTTCACCTTCGGC TCCGGCACCAAGCTGGAAATCAAG |
| CD8a Hinge | GCCCTGAGCAACAGCATCATGTACTTCAGCCACTTCGTGCCCGTGTTT CTGCCCGCCAAGCCCACCACCACCCCTGCCCCCAGACCCCCTACCCC AGCCCCCACAATCGCCAGCCAGCCCCTGAGCCTGAGGCCCGAGGCCA GCAGACCTGCCGCTGGGGGAGCCGTGCACACCAGGGGCCTGGAC |
| CD28 TM | AAGCCCTTCTGGGTGCTGGTCGTGGTCGGCGGAGTGCTGGCCTGTTA CAGCCTGCTGGTCACCGTGGCCTTCATCATCTTTTGGGTCCGCAGCAA GCGGAGCCGGCTGCTGCACAGCGACTACATGAACATGACCCCAAGG CGGCCAGGCCCCACCCGGAAGCACTACCAGCCCTATGCCCCTCCTAG GGACTTCGCCGCCTACCGGTCC |
| CD3 Zeta | AGAGTGAAGTTCAGCCGCAGCGCCGACGCCCCTGCCTACCAGCAGGG CCAGAACCAGCTGTACAACGAGCTGAACCTGGGCAGGCGGGAGGAA TACGACGTGCTGGACAAGCGCAGAGGCCGGGACCCTGAGATGGGCG GCAAGCCCAGGCGGAAGAACCCCCAGGAAGGCCTGTATAACGAACT GCAGAAAGACAAGATGGCCGAGGCCTACAGCGAGATCGGCATGAAG GGCGAGCGGCGACGCGGCAAGGGC |
| mTagBFP2 | ATGGTGTCTAAGGGCGAAGAGCTGATTAAGGAGAACATGCACATGA AGCTGTACATGGAGGGCACCGTGGACAACCATCACTTCAAGTGCACA TCCGAGGGCGAAGGCAAGCCCTACGAGGGCACCCAGACCATGAGAA TCAAGGTGGTCGAGGGCGGCCCTCTCCCCTTCGCCTTCGACATCCTGG CTACTAGCTTCCTCTACGGCAGCAAGACCTTCATCAACCACACCCAG GGCATCCCCGACTTCTTCAAGCAGTCCTTCCCTGAGGGCTTCACATGG |

| | |
|---|---|
| | GAGAGAGTCACCACATACGAAGACGGGGGCGTGCTGACCGCTACCC<br>AGGACACCAGCCTCCAGGACGGCTGCCTCATCTACAACGTCAAGATC<br>AGAGGGGTGAACTTCACATCCAACGGCCCTGTGATGCAGAAGAAAA<br>CACTCGGCTGGGAGGCCTTCACCGAGACGCTGTACCCCGCTGACGGC<br>GGCCTGGAAGGCAGAAACGACATGGCCCTGAAGCTCGTGGGCGGGA<br>GCCATCTGATCGCAAACGCCAAGACCACATATAGATCCAAGAAACCC<br>GCTAAGAACCTCAAGATGCCTGGCGTCTACTATGTGGACTACAGACT<br>GGAAAGAATCAAGGAGGCCAACAACGAGACCTACGTCGAGCAGCAC<br>GAGGTGGCAGTGGCCAGATACTGCGACCTCCCTAGCAAACTGGGGCA<br>CAAGCTTAAT |
| Complete sequence | GGATCCATGGACTGGATCTGGCGGATTCTGTTCCTGGTCGGGGCTGC<br>CACAGGCGCCCACAGCCAGGTGCAGCTGCAGCAGAGCGGCCCTGAG<br>CTGAAGAAGCCCGGCGAGACAGTCAAGATCAGCTGCAAGGCCAGCG<br>GCTACCCCTTCACCAACTACGGCATGAACTGGGTGAAACAGGCCCCA<br>GGCCAGGGACTGAAGTGGATGGGCTGGATCAACACCAGCACCGGCG<br>AGAGCACCTTCGCCGACGACTTCAAGGGCAGATTCGACTTCAGCCTG<br>GAAACCAGCGCCAACACCGCCTACCTGCAGATCAACAACCTGAAGA<br>GCGAGGACAGCGCCACCTACTTTTGCGCCAGATGGGAGGTGTACCAC<br>GGCTACGTGCCCTACTGGGGCCAGGGCACCACCGTGACCGTGTCCAG<br>CGGCGGAGGGGGCTCTGGCGGCGGAGGATCTGGGGGAGGGGGCAGC<br>GACATCCAGCTGACCCAGAGCCACAAGTTTCTGAGCACCAGCGTGGG<br>CGACCGGGTGTCCATCACCTGCAAAGCCAGCCAGGACGTGTACAACG<br>CCGTGGCCTGGTATCAGCAGAAGCCTGGCCAGAGCCCCAAGCTGCTG<br>ATCTACAGCGCCAGCAGCCGGTACACCGGCGTGCCCAGCAGGTTCAC<br>CGGCAGCGGCAGCGGCCCAGACTTCACCTTCACCATCAGCAGCGTGC<br>AGGCCGAGGACCTGGCCGTGTACTTCTGCCAGCAGCACTTCCGGACC<br>CCCTTCACCTTCGGCTCCGGCACCAAGCTGGAAATCAAGGCCCTGAG<br>CAACAGCATCATGTACTTCAGCCACTTCGTGCCCGTGTTTCTGCCCGC<br>CAAGCCCACCACCACCCCTGCCCCCAGACCCCCTACCCCAGCCCCCA<br>CAATCGCCAGCCAGCCCCTGAGCCTGAGGCCCGAGGCCAGCAGACCT<br>GCCGCTGGGGGAGCCGTGCACACCAGGGGCCTGGACAAGCCCTTCTG<br>GGTGCTGGTCGTGGTCGGCGGAGTGCTGGCCTGTTACAGCCTGCTGG<br>TCACCGTGGCCTTCATCATCTTTTGGGTCCGCAGCAAGCGGAGCCGG<br>CTGCTGCACAGCGACTACATGAACATGACCCCAAGGCGGCCAGGCCC<br>CACCCGGAAGCACTACCAGCCCTATGCCCCTCCTAGGGACTTCGCCG<br>CCTACCGGTCCAGAGTGAAGTTCAGCCGCAGCGCCGACGCCCCTGCC<br>TACCAGCAGGGCCAGAACCAGCTGTACAACGAGCTGAACCTGGGCA<br>GGCGGGAGGAATACGACGTGCTGGACAAGCGCAGAGGCCGGGACCC<br>TGAGATGGGCGGCAAGCCCAGGCGGAAGAACCCCCAGGAAGGCCTG<br>TATAACGAACTGCAGAAAGACAAGATGGCCGAGGCCTACAGCGAGA<br>TCGGCATGAAGGGCGAGCGGCGACGCGGCAAGGGCCACGACGGCCT<br>GTACCAGGGCCTGTCCACCGCCACCAAGGACACCTACGACGCCCTGC<br>ACATGCAGGCCCTGCCTCCCCGTGCTAGCGCCACGAACTTCTCTCTGT<br>TAAAGCAAGCAGGCGACGTGGAAGAAAACCCCGGTCCCATGGTGTC<br>TAAGGGCGAAGAGCTGATTAAGGAGAACATGCACATGAAGCTGTAC<br>ATGGAGGGCACCGTGGACAACCATCACTTCAAGTGCACATCCGAGGG |

| | |
|---|---|
| | CGAAGGCAAGCCCTACGAGGGCACCCAGACCATGAGAATCAAGGTG GTCGAGGGCGGCCCTCTCCCCTTCGCCTTCGACATCCTGGCTACTAGC TTCCTCTACGGCAGCAAGACCTTCATCAACCACACCCAGGGCATCCC CGACTTCTTCAAGCAGTCCTTCCCTGAGGGCTTCACATGGGAGAGAG TCACCACATACGAAGACGGGGGCGTGCTGACCGCTACCCAGGACACC AGCCTCCAGGACGGCTGCCTCATCTACAACGTCAAGATCAGAGGGGT GAACTTCACATCCAACGGCCCTGTGATGCAGAAGAAAACACTCGGCT GGGAGGCCTTCACCGAGACGCTGTACCCCGCTGACGGCGGCCTGGAA GGCAGAAACGACATGGCCCTGAAGCTCGTGGGCGGGAGCCATCTGA TCGCAAACGCCAAGACCACATATAGATCCAAGAAACCCGCTAAGAA CCTCAAGATGCCTGGCGTCTACTATGTGGACTACAGACTGGAAAGAA TCAAGGAGGCCAACAACGAGACCTACGTCGAGCAGCACGAGGTGGC AGTGGCCAGATACTGCGACCTCCCTAGCAAACTGGGGCACAAGCTTA ATGTCGAC |
